# Functional organization of social perception in the human brain

**DOI:** 10.1101/2021.12.17.473175

**Authors:** Severi Santavirta, Tomi Karjalainen, Sanaz Nazari-Farsani, Matthew Hudson, Vesa Putkinen, Kerttu Seppälä, Lihua Sun, Enrico Glerean, Jussi Hirvonen, Henry K. Karlsson, Lauri Nummenmaa

**Author notes:** **Address correspondence to**, Severi Santavirta, Turku PET Centre c/o Turku University, Kiinamyllynkatu 4-6, 20520 Turku, Finland.

## Abstract

Humans rapidly extract diverse and complex information from ongoing social interactions, but the perceptual and neural organization of the different aspects of social perception remains unresolved. We showed short film clips with rich social content to 97 healthy participants while their haemodynamic brain activity was measured with fMRI. The clips were annotated moment-to-moment for 112 social features. Cluster analysis revealed that 13 dimensions were sufficient for describing the social perceptual space. Regression analysis was used to map regional neural response profiles to different social features. Multivariate pattern analysis was then utilized to establish the spatial specificity of these responses. The results revealed a gradient in the processing of social information in the brain. Posterior temporal and occipital regions were broadly tuned to most social dimensions and the classifier revealed that these responses showed spatial specificity for social dimensions; in contrast Heschl gyri and parietal areas were also broadly associated with different social signals, yet the spatial patterns of responses did not differentiate social dimensions. Frontal and subcortical regions responded only to a limited number of social dimensions and the spatial response patterns did not differentiate social dimension. Altogether these results highlight the distributed nature of social processing in the brain.

## Introduction

Humans live in a complex and ever-changing social world, but how do we make sense of the high-dimensional and time-variable information constantly conveyed by our conspecifics? Prior functional imaging studies localized specific aspects of social perception into different brain regions (Brooks et al., 2020). Fusiform gyrus (FG) is consistently involved in the perception of faces (Haxby et al., 2000) and lateral occipitotemporal cortex (LOTC) in the perception of bodies (Downing et al., 2001). Temporoparietal junction (TPJ) is in turn involved in reflecting the mental states of others (Saxe & Kanwisher, 2003) as well as processing social context and focusing attention (Carter & Huettel, 2013). Polysensory areas in the superior temporal sulcus (STS) have been associated with multiple higher-order aspects of social perception (Deen et al., 2015; Isik et al., 2017; Lahnakoski et al., 2012; Nummenmaa & Calder, 2009), while medial frontal cortex (MFC) has been extensively studied in the context of self-representation and theory of mind (Amodio & Frith, 2006). Finally, speech-based social communication is accomplished by a network consisting of superior temporal gyrus (STG) and its proximal areas STS (Wernicke area in left pSTS), TPJ, angular gyrus, middle temporal gyrus (MTG) and inferior frontal gyrus (Broca’s area in the left IFG) (Price, 2012).

Humans can however reliably process numerous simultaneously occurring features of the social world ranging from others’ facial identities and emotions to their intentions and mental contents to the finegrained affective qualities of the social interaction. Given the computational limits of the human brain, it is unlikely that all features and dimensions of the social domain are processed by distinct areas and systems (Huth et al., 2012). Although the brain basis of perceiving specific isolated social features has been successfully delineated, the phenomenological as well as neural organization of the different social perceptual processes have remained poorly understood. Neural responses to complex stimuli cannot necessarily be predicted on statistical combination of responses to simple stimuli (Felsen & Dan, 2005). Therefor studies based on neural responses to isolated social features may not directly generalize to real-world social perception (Adolphs et al., 2016) where social features such as facial identities, body movements, and nonverbal communication often overlap with distinct temporal occurrence patterns.

In psychological domains including actions (Huth et al., 2012), language (Huth et al., 2016), and emotions (Koide-Majima et al., 2020), neuroimaging studies have tackled this issue by first generating a comprehensive set of modelled dimensions for the complex dynamic stimulus. Then, using dimension reduction techniques, they assess the representational similarities of the modelled dimensions, or the representational similarities of the brain activation patterns associated with each dimension. For example, a recent study found that linguistic and visual semantic representations converge so that visual representations locate on the border of occipital cortex and that linguistic representations are located anterior to the visual representations (Popham et al., 2021). However, a detailed representational space for social features at both perceptual and neural level is currently lacking.

We define social perception as perception of all possible information relevant to interpret social interaction. To our knowledge, there is no consensus on a combined taxonomy for this broad definition. In social psychology, social situation has been described as a triad of person, situation and consequent behaviour (Lewin, 1936) where these elements have close interact between each other (Funder, 2006). However, data-driven taxonomies have only been proposed for the elements separately. Person perception has been extensively studied and person characteristics can be categorised as a limited set of trait dimensions, such as Big Five (Goldberg, 1990) or Big Six (Lee & Ashton, 2004). For psychological situations, data-driven lexical studies have proposed limited dimensionality (Parrigon et al., 2017; Rauthmann et al., 2014). Recently, in behavioural domain categorization of human actions have also been proposed (Thornton & Tamir, 2022). For two reasons, these established taxonomies are suboptimal for studying social perception as whole. First, these taxonomical studies base their results on questionnaires regarding social situations or rated similarities of different words describing social situations instead of the actual perception of social situations in real-life dynamic environment. Second, since the three elements have a close interaction, it would be sensible to study them together. Therefor our approach involves first collecting ratings of a large set of perceived social features from the stimulus used in this neuroimaging study and then limiting the social perceptual space with clustering analysis.

Commonly applied univariate analyses modeling the BOLD response in each voxel or region separately cannot reveal the specificity of spatial brain activation patterns resulting from the perception of different social features. Consequently, they do not allow testing whether different perceptual features can be reliably discriminated based on their spatial brain activation patterns. Multivariate pattern analysis (MVPA) allows the analysis of information carried by fine-grained spatial patterns of brain activation (Tong & Pratte, 2012). Pattern recognition studies have established that regional multivariate patterns allow distinguishing brain activation related to multiple high-level social features such as faces (Haxby et al., 2001) and their racial group (Brosch et al., 2013) in FG and facial expressions in FG and STS (Harry et al., 2013; Said et al., 2010; Wegrzyn et al., 2015). Perception of different goal-oriented motor actions with different levels of abstraction can be decoded in LOTC and in inferior parietal lobe, suggesting that these regions process the abstract concepts of the goal-oriented actions, not just their low-level visual properties (Wurm & Lingnau, 2015). Furthermore, decoding of goal-oriented actions was successful in LOTC when subjects observed the actions in both first and third person perspectives (Oosterhof et al., 2012). It however remains unresolved how specific these regional response profiles are across different social perceptual features.

### The current study

In this fMRI study, we mapped the perceptual and neural representations of naturalistic social episodes using both univariate and multivariate analyses. We used short film clips as stimuli because cinema contains rich and complex social scenarios and as it also elicits strong and consistent neural responses in functional imaging studies (Hasson et al., 2010; Lahnakoski et al., 2012). We first aimed at establishing a perception-based taxonomy of the social dimensions that human observers use for describing social scenarios, and then mapping the brain basis of this social perceptual space. We mapped the perceptual space of social processes based on subjective annotations of a large array of social features (n=112) in the films (n=96). We then used dimension reduction techniques to establish the representational space of social perception, and to reduce the multidimensional space into a limited set of reliable perceptual dimensions of social features. Using combination of univariate regression analysis, and multivariate pattern analysis we established that posterior temporal and occipital regions are the main hubs for social perception and that brain shows a gradient in social perceptual processing from broadly tuned but spatially dimension-specific responses in posterior temporal and occipital regions towards more selective responses in frontal and subcortical areas.

## Materials and methods

### Participants

Altogether 102 volunteers participated in the study. The exclusion criteria included a history of neurological or psychiatric disorders, alcohol or substance abuse, BMI under 20 or over 30, current use of medication affecting the central nervous system and the standard MRI exclusion criteria. Two additional subjects were scanned but excluded from further analyses because unusable MRI data due to gradient coil malfunction. Two subjects were excluded because of anatomical abnormalities in structural MRI and additional three subjects were excluded due to visible motion artefacts in preprocessed functional neuroimaging data. This yielded a final sample of 97 subjects (50 females, mean age of 31 years, range 20 – 57 years). All subjects gave an informed, written consent and were compensated for their participation. The ethics board of the Hospital District of Southwest Finland had approved the protocol and the study was conducted in accordance with the Declaration of Helsinki.

### Stimulus

To map brain responses to different social features, we used our previously validated socioemotional “localizer” paradigm that allows reliable mapping of various social and emotional functions (Karjalainen et al., 2017; Karjalainen et al., 2019; Lahnakoski et al., 2012; Nummenmaa et al., 2021). The original experiment using this stimulus described the experimental design and stimulus selection in detail (Lahnakoski et al., 2012). Briefly, the subjects viewed a medley of 96 movie clips (median duration 11.2 seconds, range 5.3 – 28.2 seconds, total duration 19 min 44 seconds) that have been curated to contain large variability of social and emotional content. The videos were extracted from mainstream Hollywood movies with audio track in English. To limit experiment duration 87 of the previously validated 137 clips were selected. 71 of these clips contained people in various social situations and contexts (one person: 15, two people: 22, more than two people: 34). To distinguish person perception from other audiovisual perception the stimulus contained four clips with animals and 12 control clips without people (showing e.g. scenery and objects). Additionally, nine erotic scenes showing heterosexual intercourse were added to broaden the emotional content of the original stimulus. Short descriptions about movie clips can be found from **Table SI-1**. Because this task was designed to map neural processing of naturalistic socioemotional events, the clips were not deliberately matched with respect to, for example, human motion or optic flow. The videos were presented in fixed order across the subjects without breaks. Subjects were instructed to view the movies similarly as if they were viewing a film at a cinema or at home and no specific task was assigned. Visual stimuli were presented with NordicNeuroLab VisualSystem binocular display. Sound was conveyed with Sensimetrics S14 insert earphones. Stimulation was controlled with Presentation software. Before the functional run, sound intensity was adjusted for each subject so that it could be heard over the gradient noise.

### Neuroimaging data acquisition and preprocessing

MR imaging was conducted at Turku PET Centre. The MRI data were acquired using a Phillips Ingenuity TF PET/MR 3-T whole-body scanner. High-resolution structural images were obtained with a T1-weighted (T1w) sequence (1 mm^3^ resolution, TR 9.8 ms, TE 4.6 ms, flip angle 7°, 250 mm FOV, 256 × 256 reconstruction matrix). A total of 467 functional volumes were acquired for the experiment with a T2*-weighted echo-planar imaging sequence sensitive to the blood-oxygen-level-dependent (BOLD) signal contrast (TR 2600 ms, TE 30 ms, 75° flip angle, 240 mm FOV, 80 × 80 reconstruction matrix, 62.5 kHz bandwidth, 3.0 mm slice thickness, 45 interleaved axial slices acquired in ascending order without gaps).

The functional imaging data were preprocessed with FMRIPREP (Esteban et al., 2019) (v1.3.0), a Nipype (Gorgolewski et al., 2011) based tool that internally uses Nilearn (Abraham et al., 2014). During the preprocessing, each T1w volume was corrected for intensity non-uniformity using N4BiasFieldCorrection (v2.1.0) (Tustison et al., 2010) and skull-stripped using antsBrainExtraction.sh (v2.1.0) using the OASIS template. Brain surfaces were reconstructed using recon-all from FreeSurfer (v6.0.1) (Dale et al., 1999), and the brain mask estimated previously was refined with a custom variation of the method to reconcile ANTs-derived and FreeSurfer-derived segmentations of the cortical greymatter of Mindboggle (Klein et al., 2017). Spatial normalization to the ICBM 152 Nonlinear Asymmetrical template version 2009c (Fonov et al., 2009) was performed through nonlinear registration with the antsRegistration (ANTs v2.1.0) (Avants et al., 2008), using brain-extracted versions of both T1w volume and template. Brain tissue segmentation of cerebrospinal fluid, white-matter and grey-matter was performed on the brain-extracted T1w image using FAST (Zhang et al., 2001) (FSL v5.0.9).

Functional data were slice-time-corrected using 3dTshift from AFNI (Cox, 1996) (v16.2.07) and motion-corrected using MCFLIRT (Jenkinson et al., 2002) (FSL v5.0.9). These steps were followed by coregistration to the T1w image using boundary-based registration (Greve & Fischl, 2009) with six degrees of freedom, using bbregister (FreeSurfer v6.0.1). The transformations from motion-correction, coregistration, and spatial normalization were concatenated and applied in a single step using antsApplyTransforms (ANTs v2.1.0) using Lanczos interpolation. Independent-component-analysis-based Automatic Removal Of Motion Artifacts (ICA-AROMA) was used to denoise the data nonaggressively after spatial smoothing with 6-mm Gaussian kernel (Pruim et al., 2015). The data were then detrended using 240-s-Savitzky–Golay filtering to remove the scanner drift (Cukur et al., 2013), and finally downsampled to original 3 mm isotropic voxel size. The BOLD signals were demeaned to make the regression coefficients comparable across different individuals (Chen et al., 2017). First and last two functional volumes were discarded to ensure equilibrium effects and to exclude the time points after the stimulus had ended.

### Stimulus features

Five individuals not participating in fMRI rated the 112 predefined social features (see **Table SI-2**) from the film clips. We selected a broad range of socioemotional features describing persons, social situations and behaviours from following categories: sensory input (e.g. smelling, tasting), basic body functions (e.g. facial expressions, walking, eating), person characteristics (e.g. pleasantness, trustworthiness) and person’s inner states (e.g. pleasant feeling, arousal), social interaction signals (e.g. talking, communicating with gestures) and social interaction characteristics (e.g. hostility, sexuality). Collecting perceptual ratings from a large set of individual social features enables reliable mapping of the whole social perceptual space that can be derived from the stimulus film clips and ensures that the data-driven dimensionality arises from the used stimulus. It was stressed to the observers that they should rate the perceived features of the social interaction rather than the observer’s own inner states (such as emotions evoked by the films). The ratings were collected separately for each video clip in short time intervals (median 4.0 sec, range: 3.1 — 7.3 seconds). Features were annotated in a continuous and abstract scale from “absent” to “extremely much”. For analyses the ratings were transformed to continuous scale from 0 (absent) to 100 (extremely much). Annotators watched the video clips altogether 12 times, rating an average of 10 features on each viewing to reduce the cognitive load. The ratings were done using an online rating platform Onni (http://onni.utu.fi) developed at Turku PET Centre (Heikkilä et al., 2020).

### Feature reliability

We first evaluated whether the *a priori* features were frequently and consistently perceived in the stimulus films. Features with low occurrence rate and/or inter-rater reliability were excluded, because i) high occurrence rate is needed to reliably estimate the stimulus-dependent variation in BOLD signal, and ii) high inter-rater reliability is necessary to study brain activity in a sample of participants independent from the raters. The occurrence rate was defined as the number of time points where the mean (minus standard error of the mean) of the rating exceeded 5 (on a scale ranging from 0 to 100). Features were included in the analyses if they occurred at least five times throughout the experiment; this was estimated to yield sufficient statistical power in the BOLD-fMRI GLM analyses. Inter-rater reliability of the features was assessed using intra-class correlation coefficient (ICC) as calculated in the R package psych (https://cran.r-project.org/package=psych). ICC(A,1) was selected as appropriate model for ICC since it treats both video clips and raters as random effects and measures the absolute agreement between raters (McGraw & Wong, 1996). ICCs below 0.5 are considered poor (Koo & Li, 2016), and we thus only included features with ICC over 0.5. 45 features satisfied both criteria. The occurrence rate and interrater reliability of each feature are shown in **Figure SI-1**.

### Dimension reduction

The reliable 45 features were linearly correlated (**Figure 1**) and it is unlikely that each social feature is processed in different brain regions or networks. We performed dimension reduction with hierarchical clustering on the correlation matrix of selected features to define the perceptual dimensions that characterize different aspects of social interaction. Clustering was chosen over principal component analysis for easier interpretation of the dimensions, and as this allowed us to model also features or their combinations which may be salient and reliable but not necessarily share a large proportion of variance with other variables. Initially we chose Pearson correlation as the similarity measure because the cooccurrence of features measured in abstract and possibly not strictly continuous scale is more interesting than the absolute distance between them (considered in PCA). Unweighted pair group method with arithmetic mean (UPGMA), as implemented in R, was used as the clustering algorithm (https://www.rdocumentation.org/packages/stats/versions/3.6.2/topics/hclust). Other average linkage clustering methods implemented in the R package (WPGMA, WPGMC and UPGMC) yielded highly similar clustering hierarchy. Hierarchical clustering requires a desired number of resulting clusters as an input for automatic definition of cluster boundaries from hierarchical tree (**Figure SI-2**). To estimate the optimal number of clusters we chose three criteria that the clustering result should satisfy. These were cluster stability, theoretically meaningful clustering, and sufficient reduction in collinearity between the clusters. To assess the stability of clusters we conducted a consensus clustering analysis with ConsensusClusterPlus R package (Wilkerson & Hayes, 2010). Theoretically meaningful clustering was then assessed, and collinearity was measured using Pearson correlation and variance inflating factor (VIF). Detailed information of the cluster analysis and consensus clustering results can be found in Supplementary Materials (see also **Figure SI-3**). Cluster analysis grouped social features into six clusters and seven independent features not belonging to any cluster (**Figure 1**) and these social dimensions formed the final model for social perception. The cluster regressors were created by averaging across the individual feature values in each cluster (**Figure SI-4**).

**Figure 1.**
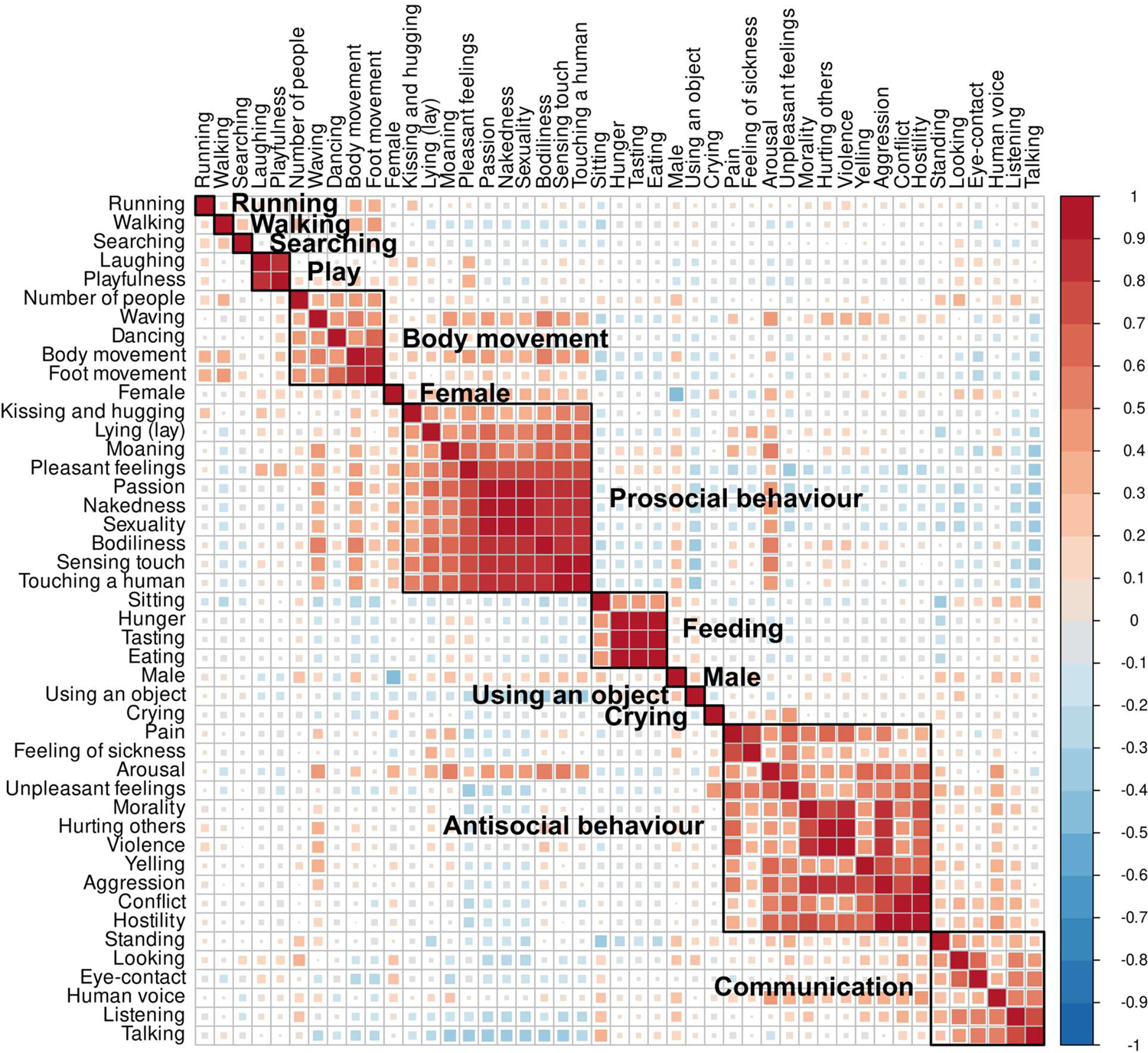
The results of hierarchical clustering of reliably rated social features. The correlation matrix is ordered hierarchically, and clustering results (k=13) are shown. Hierarchical clustering analysis identified that social perceptual space of the stimulus can be reduced to six clusters and seven individual features.

### Modelling low-level sensory features

Our goal was to map perceived social dimensions in the human brain. The stimulus film clips were not balanced by their low-level audiovisual properties thus these were controlled statistically when estimating the unique contribution of social dimensions to the BOLD signal. We extracted 14 different dynamic audiovisual properties from the stimulus film clips including six visual features (luminance, first derivative of luminance, optic flow, differential energy, and spatial energy with two different frequency filters) and eight auditory features (RMS energy, first derivative of RMS energy, zero crossing, spectral centroid, spectral entropy, high frequency energy and roughness). Optic flow was estimated with opticalFlowLK - function with basic options (https://www.mathworks.com/help/vision/ref/opticalflowlk.html). Custom functions were used for estimating other visual features (see Code availability). Auditory features were extracted using MIRToolbox1.8.1 (Olivier Lartillot & Toiviainen, 2007). First eight principal components (PCs) explaining over 90 % of the total variance were selected as regressors for low-level audiovisual features. As the stimulus film clips included control clips with no human interaction, we created a “nonsocial” block regressor by assigning a value of 1 to the time points where the stimulus did not contain people, human voice, or animals. A low-level model was formed by combining the eight audiovisual PCs, the nonsocial regressor and subjectwise mean signals from cerebrospinal fluid (CSF) and white matter (WM). See **Figure SI-5** for correlations between low-level features and social dimensions.

### Modelling brain responses to social perceptual dimensions

Ridge regression (Hoerl & Kennard, 1970) was used to estimate the contributions of the low-level features and cluster-based composite social dimensions to the BOLD signals for each subject. Ridge regression was preferred over ordinary least squares (OLS) regression because even after dimension reduction, the social regressors were moderately correlated (range: −0.38 – 0.32) and we wanted to include all perceptual dimensions in the same model to estimate their unique contributions to the BOLD signals. We also wanted to avoid overfitting while retaining generalizability of the results. To conservatively control for low-level features, the demeaned BOLD signals were first predicted with the low-level model and the residual BOLD signals were then used as input in the following regression analysis with the social stimulus model. The low-level regressors were still included as nuisance covariates in the analysis of social dimensions for the possible interaction between the social dimensions and low-level features. In both consecutive analyses ridge parameter was optimized using leave-one-subject-out cross-validation. Prior to statistical modelling the regressor time series were convolved with the canonical HRF and the columns of the design matrices were standardised (*μ* = 0, *σ* = 1). Detailed description of ridge regression modelling is included in supplementary materials (**Figure SI-6**).

Ridge regularization parameter optimisation is computationally prohibitive when considering all the voxels in the brain since the leave-one-out cross-validation loads the complete voxelwise data into memory requiring constantly 70-80 Gb of memory even when considering 20 % of the voxels in the brain. Optimizing ridge penalty for each voxel separately could have yielded in large differences in the penalty parameters values throughout the brain thus making it more difficult to interpret the regional differences in the results. Thus, we selected an unbiased sample of grey matter voxels for the optimisation by randomly sampling 20 % of grey matter voxels uniformly throughout the brain. Only voxels within population level EPI mask where the population level probability of grey matter was over 0.5 were available for sampling. Consequently, a uniformly distributed sample of ~5000 voxels was selected for ridge parameter optimization.

For the social perceptual model, the regression analysis was run both at voxel-level and at region-of-interest (ROI) level. The population level EPI-mask was used in all analyses to include only voxels with BOLD signal from each subject and thus brain areas including parts of orbitofrontal, inferior temporal and occipital pole areas were not included in the analyses. In voxel-level analysis, subject-level ß-coefficient-maps were subjected to group-level analysis to identify the brain regions where the association between intensity of each social dimension and haemodynamic activity was consistent across the subjects. Voxels outside the population level EPI mask were excluded from the analysis. Statistical significance was identified using the randomise function of FSL (Winkler et al., 2014). Voxel-level FDR with q-value of 0.05 was used to correct for multiple comparisons (Benjamini & Hochberg, 1995). Anatomical ROIs were extracted from AAL2 atlas (Rolls et al., 2015). ROIs, where at least 50% of voxels were outside the population level EPI mask, were excluded from the analysis and only voxels within population level EPI mask were considered for the included ROIs. This resulted in inclusion of 41 bilateral ROIs into the analysis. A parametric T-test on the ß-weights of a ROI was used to assess statistical inference across subjects. ROI-analysis results were considered significant with P-value threshold of 0.05 Bonferroni corrected for multiple comparisons. The results for ROI analyses are reported as union of bilateral ROIs.

### Multivariate pattern analysis of social perceptual dimensions

To reveal the regional specialization in processing of different social features, between-subject classification of 11 perceptual dimensions^1^ was performed in Python using the PyMVPA toolbox (Hanke et al., 2009). The aim of the classification analysis was to complement univariate regression analysis by testing whether the human brain expressed regional specificity for distinct social dimensions. This approach was based on classification of discrete social dimensions from brain activity, rather than computationally more complex approach to predict actual values of multiple social predictors simultaneously based on brain activity. For this kind of classification, only one dimension label for each time point (each TR) could be given. In continuous signal, the choice of the best label for each time point was not always obvious, because more than one social feature could be present simultaneously and is likely that unusual or rarely present social information draws more attention (“Somebody starts crying”) than constantly present information (“People are talking”). To resolve this issue, we first normalized the dimension rating time series (*μ* = 0, *σ* = 1) and then, for each time point, chose the feature with the highest Z-score as the category label for that time point. To ensure that the included time points would be representative of the assigned categories, we chose only time points where Z-scores for the chosen dimension were positive. With this procedure we ensured that each time point holds a label of a representative category and that rarely present information is weighted more than constantly present information.

Classifying every time point separately is computationally prohibitive and single EPI scans are noisy. Moreover, it cannot be assumed that adjacent time points assigned with the same label would be independent from each other. Accordingly, we split the data into 29 time windows and all time points with the same label within a time window were considered as a single event of that class. The number of time windows was selected based on the fact that the response length of canonical HRF is approximately 30 seconds and therefor over 30 second time windows would be less dependent from each other than shorter time windows while the data would contain enough events for classification. The time window boundaries were adjusted so that adjacent time points with the same label would not be interspersed to different time windows because temporal autocorrelation of adjacent time points may yield in artificial increase of the classification accuracy. After adjustment, the average time window length was 39 seconds (range: 34 sec – 49 sec). The time windows were longer than the movie clips and therefor timepoint from different clips with similar social context could be judged as one event if they belong to same time window. Altogether the data consisted of 87 events for each subject. These events include Using an object: 16, Communication 15, Antisocial behaviour: 11, Feeding: 10, Walking: 9, Prosocial behaviour: 8, Body movement: 5, Crying: 4, Play: 4, Running: 3 and Searching: 2.

An ordinary least squares GLM was fit for the residual BOLD time series (after regressing out the low-level features) in each time window and social dimension and the resulting subjectwise ß-images were used as input for the multivariate pattern analysis (MVPA). The input residual BOLD time series were normalized (*μ* = 0, *σ* = 1) before application of the GLM. A neural network (NN) model (https://scikit-learn.org/stable/modules/generated/sklearn.neural_network.MLPClassifier.html) was trained to classify the perceptual dimensions using leave-one-subject-out cross-validation, where the model was trained on the data from all except one subject and tested on the hold-out subject’s data; this procedure was repeated N times so that each subject was used once as the hold-out subject. Such leave-one-subject-out crossvalidation tests the generalizability of the results across the sample of the subjects. The analysis was performed using whole brain data (with non-brain voxels masked out) and regional data using anatomical ROIs. In the whole-brain analysis, an ANOVA feature selection was applied to the training set within each cross-validation and 3000 voxels with the highest F-score were selected. The regional MVPA was first performed using data form all voxels within a region. To control for the effect of ROI size to the classification accuracy the regional MVPA was also performed with an ANOVA feature selection where the size of the smallest ROI (lateral orbitofrontal cortex, 119 voxels) was selected as the number of features for the feature selection.

Hyperparameters of the NN algorithm were optimized within a limited set of predefined hyperparameter values in the whole brain analysis. Hyperparameter values reflecting the best prediction accuracy with acceptable runtime were used in both full brain and ROI analyses (see **Table SI-3** for hyperparameter tuning). The optimized NN included two hidden layers with 100 nodes in each (alpha = 1.00, max_iter 500, other hyperparameters set to default). In the model learning process, the order of events was shuffled in each training iteration which minimized the model’s ability to learn the order of the events in the stimulus. A support vector machine (SVM) classifier had similar classification accuracy in the whole brain analysis, but NN model was chosen because the computation time was shorter and the variance of classification accuracies between subjects were lower with NN model compared to SVM classifier.

Classification accuracy was quantified by computing the proportion of correctly classified events relative to the total number of events (i.e., recall). To estimate the null distribution, the following procedure was repeated 500 times: we 1) randomly shuffled social class labels; 2) ran the whole-brain MVPA with 97 leave-one-subject-out cross-validations, where the classifier was trained on the data with shuffled labels from N-1 subjects and tested on data with correct labels from the remaining subject; and 3) calculated the classification accuracies on each of the 500 iterations. The null distribution estimation was computationally prohibitive as one iteration took approximately one hour, and we decided that 500 iterations would be sufficient to assess the statistical significance of our findings. If the true accuracy was larger than 99% of the accuracies obtained with the randomly shuffled labels, the true accuracy was considered significant with an alpha of 0.01. We cannot assume that the null distribution of classification accuracies for each class is equal and center around the naive chance level because the number of events is unbalanced between classes. For this reason, we only report if the total accuracy of the classification is statistically significant. In the whole-brain classification we also report the precision of the classifications which is the number of correct predictions for a class divided by the total number of predictions into that class. In ROI analyses, the statistical differences between regional classification accuracies were tested using paired T-tests between subjectwise classification accuracies between each pair of regions.

### Inter-subject correlation analysis

Watching films synchronizes brain activity between different individuals particularly in the occipital, temporal, and parietal regions of the brain and the synchronization of brain activity can be measured with inter-subject correlation (ISC) analysis (Hasson et al., 2004). As the only variable factor in the experiment is the time-varying audiovisual stimulus, ISC analysis captures the shared stimulus-dependent activation in the brain. It is well known that ISC is greatest on the sensory cortices, but an important yet unresolved question is which variables drive the degree of synchronization of BOLD response. Some prior studies suggest that emotions and top-down perspectives play a role (Lahnakoski et al., 2014; Nummenmaa et al., 2014), but the role of social features remain unknown. As a *post hoc* analysis, we assessed whether the regional differences in brain response profiles for social dimensions relate to the inter-subject response reliability of BOLD response. To this end, we calculated the ISC across subjects over the whole experiment and compared the regional ISC with the results from regression and MVPA analyses. ISC-toolbox with default settings was used for ISC calculations (Kauppi et al., 2014).

## Results

### How people perceive the social world?

A total of 45 out of the 112 social features had sufficient inter-rater reliability and occurrence rate (see **Figure SI-1**). Hierarchical clustering identified six clusters that were labelled as “Antisocial behaviour”, “Prosocial behaviour”, “Communication”, “Body movement”, “Feeding” and “Play”. Seven perceptual dimensions did not link with any cluster and were analysed separately. These dimensions were “Using an object”, “Crying”, “Male”, “Female”, “Running”, “Walking” and “Searching”. **Figure 1** shows the clustering of the dimensions. Median pairwise correlation between any two of the 13 dimensions was 0.02 (range: −0.38 — 0.32) and the maximum variance inflation factor (VIF) in the design matrix excluding nuisance covariates was 3.3 (male regressor) and median VIF value was 1.3. These diagnostics indicate that regression coefficients for the dimensions will be stable in linear model estimations and, and they could thus be included in the same model. See **Figure SI-4** for visualized time series of social dimensions and **Figure SI-5** for correlations matrices for low-level features and social dimensions.

### Cerebral topography of social perception

Regularized ridge regression was used to establish the full-volume activation patterns for 13 perceptual social dimensions (**Figure 2**). Social information processing engaged all brain lobes and both cortical and subcortical regions. Robust responses were observed in occipital, temporal, and parietal cortices (**Figure 3**). There was a clear gradient in the responses, such that posterior temporal, occipital and parietal regions showed the strongest positive association with most of the social dimensions, with significantly less consistent activations in the frontal lobes and subcortical regions. Yet, frontal, and subcortical activations were also observed for some dimensions such as prosocial behaviour, antisocial behaviour and feeding.

**Figure 2.**
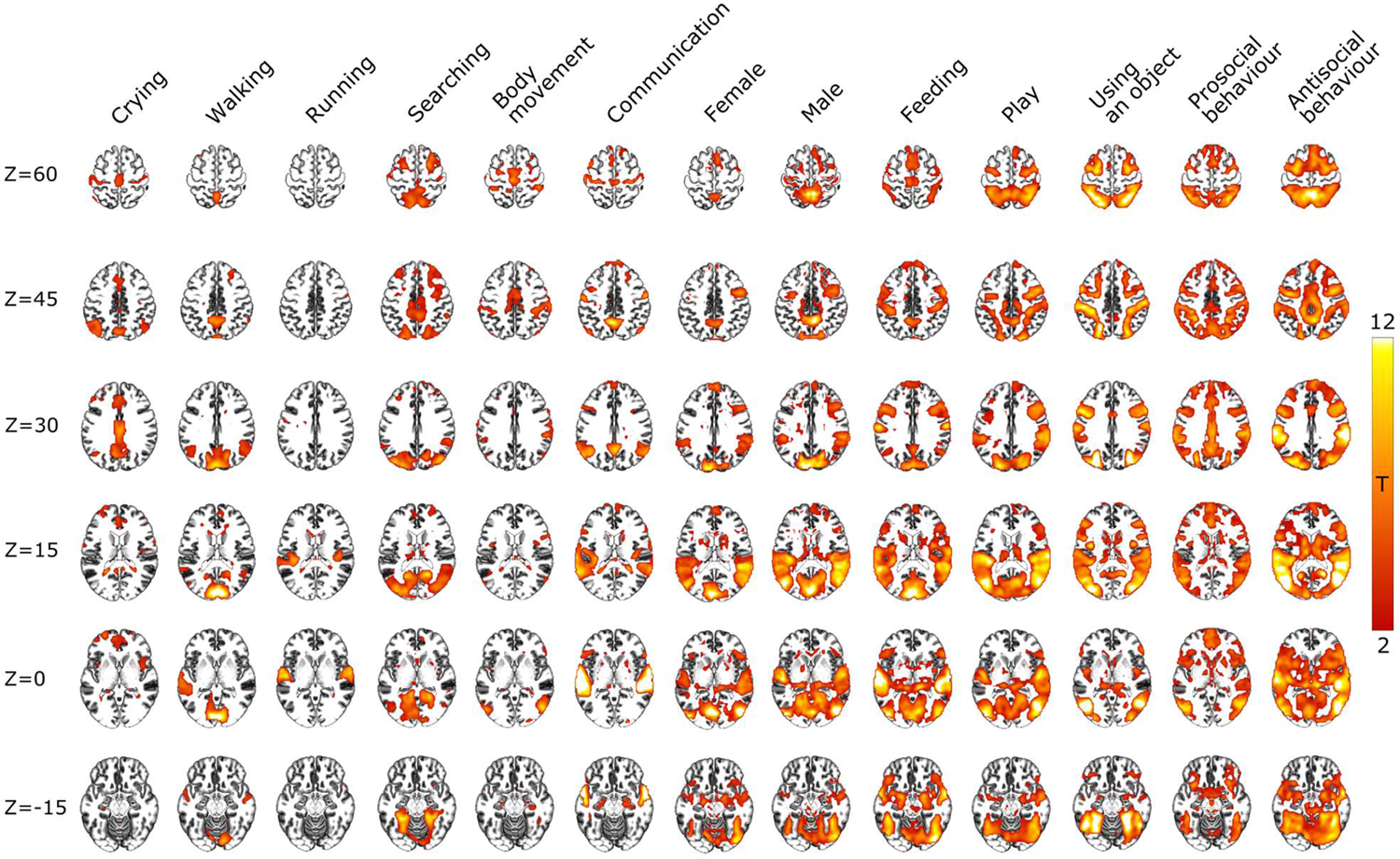
Brain regions showing increased BOLD activity for the social dimensions. Results show the voxelwise T-values (FDR-corrected, q = 0.05) of increased BOLD activity for each social dimension from the multiple regression analysis. The results are also visualized as cortical inflation in **Figure SI-7**.

**Figure 3.**
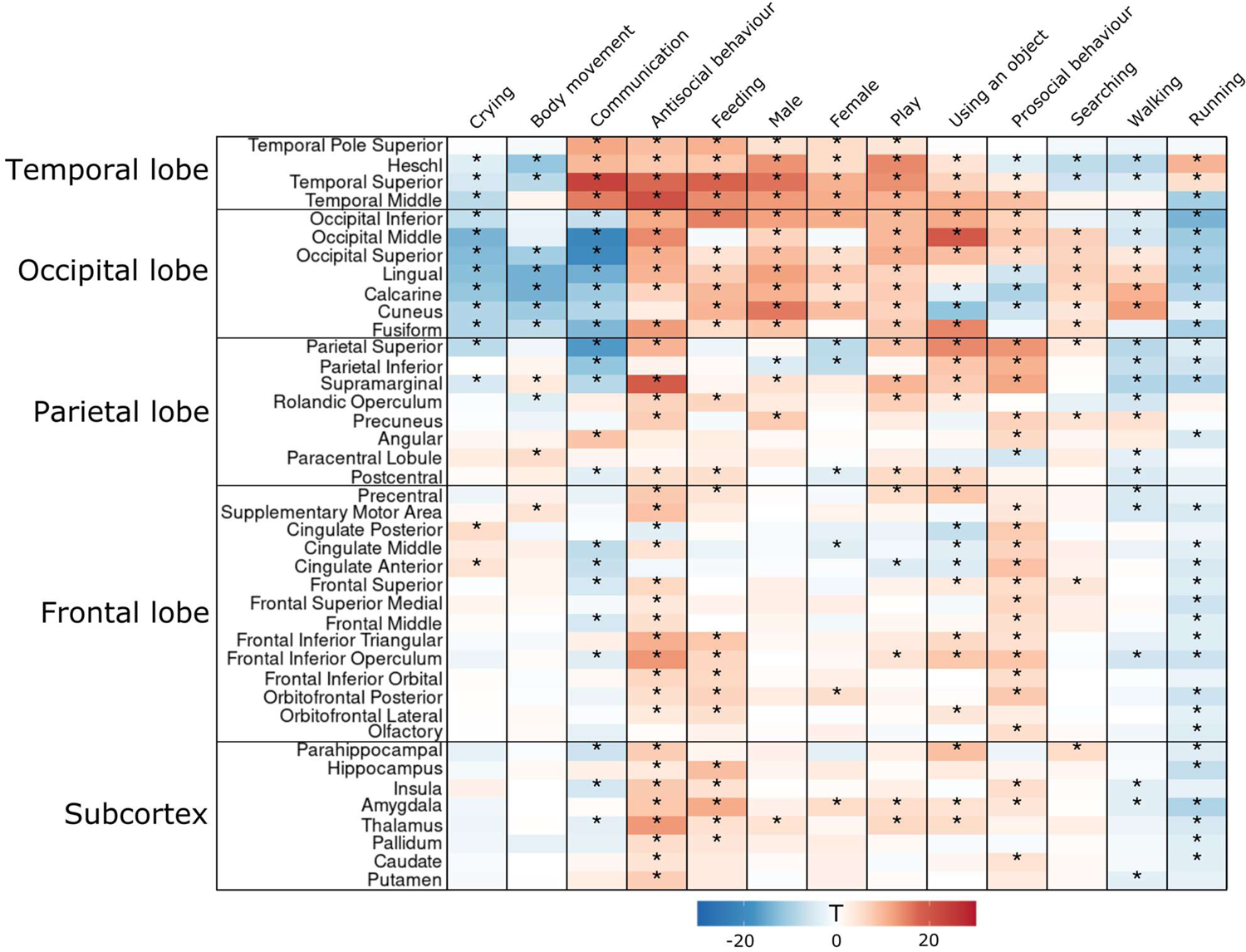
Regional results from the multiple regression analysis. A heatmap shows T-values for regression coefficients in each ROI and perceptual dimension. Statistically significant (p < 0.05, Bonferroni-corrected for each dimension independently) ROIs are marked with an asterisk.

In ROI analysis, broad responses for social dimensions were observed in STG and MTG with strongest responses for communication and antisocial behaviour, respectively. In parietal lobe, all regions except angular gyrus and paracentral lobule associated with a wide range of perceptual dimensions. In frontal regions the associations between social dimensions and haemodynamic activity were less consistent than in more posterior regions, yet still statistically significant in some of the regions including IFG, cingulate cortex and precentral gyrus. Most consistent frontal effects were found for prosocial and antisocial behaviour. For subcortical regions the observed associations were generally weak. Most notable subcortical associations with perceptual dimensions were seen in amygdala and thalamus. Consistent negative associations were restricted to occipital lobe and were observed for communication, crying, body movement and running.

### Brain gradient of social perception

**Figure 4a** shows the cumulative brain activation maps for all 13 perceptual dimensions. There was a gradient in the regional selectivity for social dimensions. Posterior temporal and occipital cortices as well as parietal cortices responded to most social dimensions, while responses become more selective in the frontal cortex although IFG, precentral gyrus and the frontal part of the medial superior frontal gyrus (SFG) had some consitency in their response profiles. Because the same stimulus was used across the subjects, we hypothesized that the brain activation in the areas with the broadest response profiles would be temporally most synchronized across subjects. We thus calculated the ISC of brain activation over the whole experiment (**Figure 4b**) and correlated the regional ISC values with corresponding response selectivity values (i.e. number of social features resulting in significant activations in each region). Scatterplot in **Figure 4c** shows the association between ISC and corresponding brain response selectivity for perceptual dimensions (Pearson r = 0.86).

**Figure 4.**
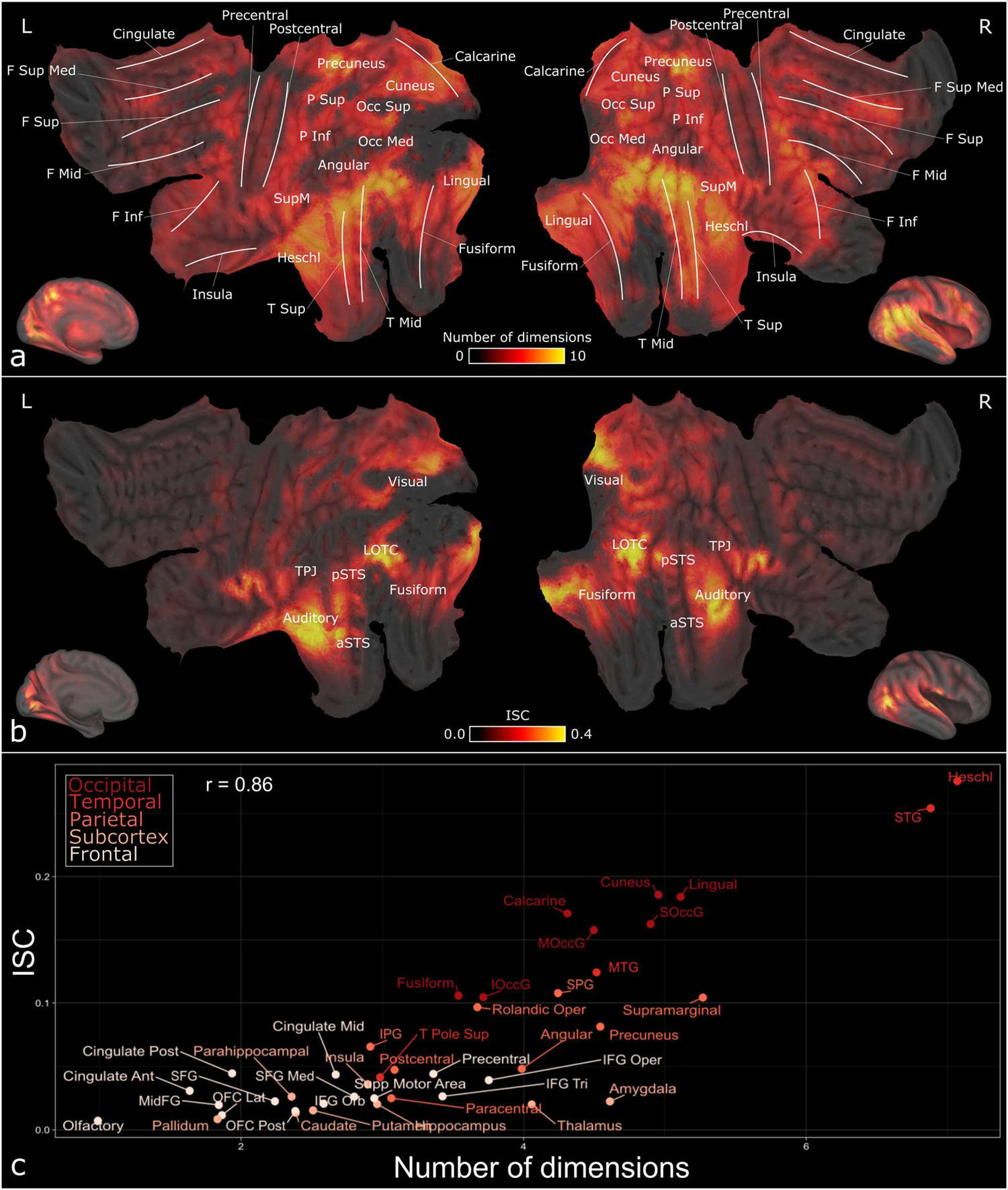
(a) Cumulative activation map for social dimensions. Voxel intensities indicate how many social dimensions (out of 13) activated the voxel statistically significantly (FDR-corrected, q = 0.05). White lines indicate the localizations of major gyri. (b) Significant ISC (FDR-corrected, q = 0.05) across subjects over the whole experiment (c) Scatterplot showing the association between ISC and tuning for social perceptual dimensions. ISC is plotted in Y-axis and the X-axis shows how many social dimensions (out of 13) associated significantly with BOLD response. Regional values are calculated as the average over all regional voxels. CARET software (Van Essen, 2012) was used for mapping results from ICBM 152 Nonlinear Asymmetrical template version 2009c space to the flatmap surface.

### Multivariate pattern analysis

Finally, we trained a between-subject neural network model to decode presence of perceptual social dimensions from the spatial haemodynamic activation patterns to reveal which social dimensions are consistently represented in each cerebral region. Whole brain classification was performed in 3000 voxels that passed through the ANOVA feature selection. Most of the selected voxels (**Figure 5a**) localized into temporal (STG, MTG, Heschl gyrus and superior temporal pole), occipital (calcarine and lingual gyri, cuneus, FG, superior occipital gyrus (SOccG), middle occipital gyrus (MoccG) and inferior occipital gyrus (IoccG)) and parietal cortices (supramarginal, superior parietal gyrus (SPG) and inferior parietal gyrus (IPG)). The permuted chance level for the total classification accuracy in the whole brain analysis was 0.128 which is above naïve chance level 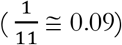. At the whole brain level, the NN model was able to classify all 11 social dimensions significantly above chance level with the total classification accuracy of 0.52 (p<0.01). Classification accuracies/precisions for each social dimension were: walking: 0.49/0.51, using an object: 0.53/0.50, searching: 0.70/0.69, running 0.56/0.62, prosocial behaviour 0.45/0.48, play 0.53/0.51, feeding 0.46/0.48, crying 0.46/0.51, communication 0.55/0.55, body movement 0.52/0.50 and antisocial behaviour 0.55/0.53 (**Figure 5a**).

**Figure 5.**
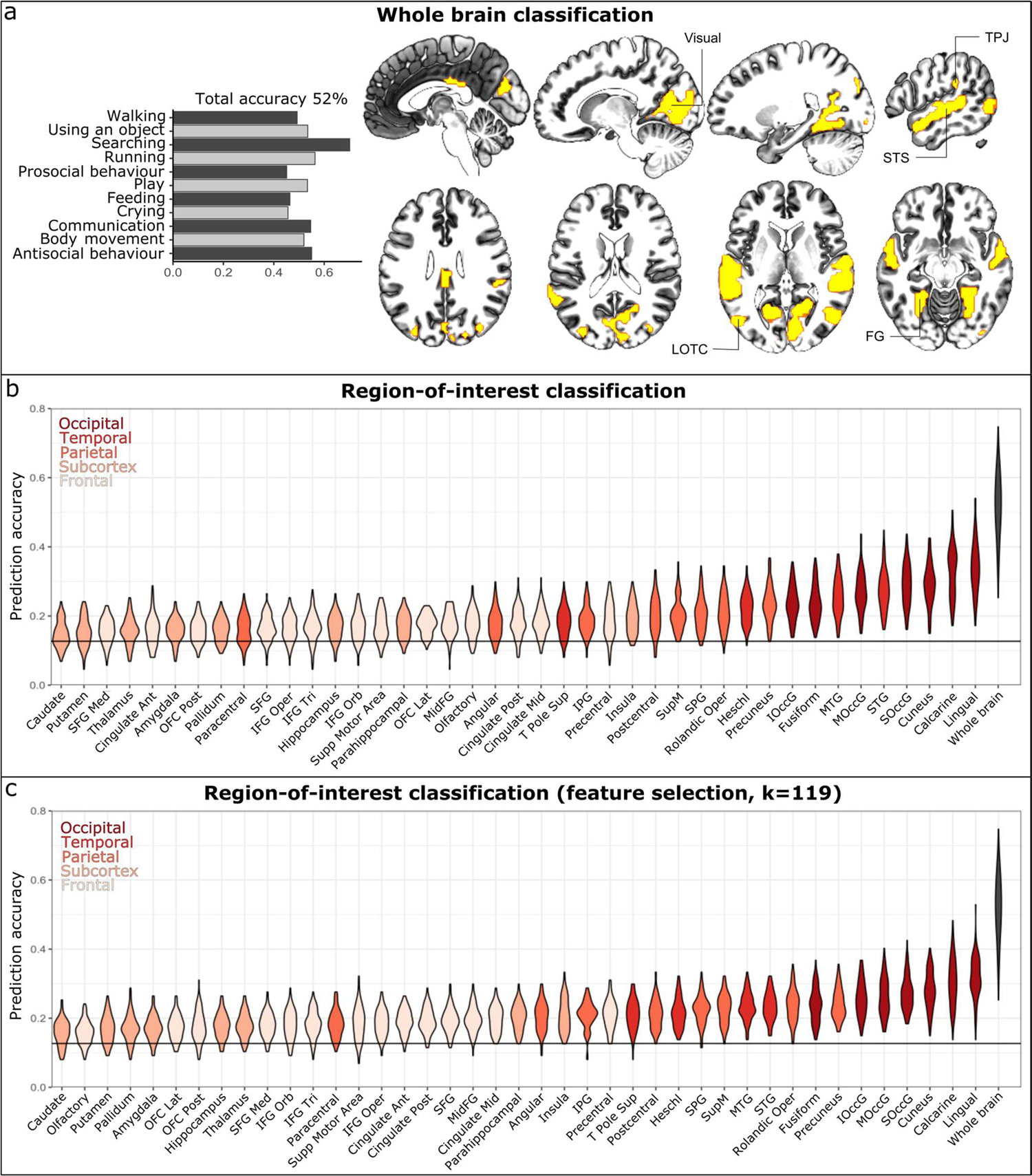
Results from the multivariate pattern analysis of social dimensions. (a) Whole brain classification accuracies and the localization of ANOVA selected voxels used in the whole brain classification analysis. (b) Regional classification accuracies compared with the whole brain classification accuracy (black). The permuted chance level accuracy (acc = 0.128) is shown as a horizontal line. The mean prediction accuracy was significantly (p<0.01) above the chance level accuracy in the whole brain analysis and for each region-of-interest. (c) Regional classification accuracies using only 119 voxels (the size of the smallest region) with the highest F-scores as input for the classifier.

The classification was also performed within anatomical ROIs (**Figure 5b**). Most accurate classifier performance was observed in lingual gyrus (0.34, p<0.01), calcarine gyrus (0.33, p<0.01), cuneus (0.29, p<0.01), SOccG (0.29, p<0.01), MOccG (0.27, p<0.01), STG (0.27, p<0.01) and MTG (0.25, p<0.01). Although the prediction accuracies were statistically significantly above permuted chance level for each ROI, the gradient in brain responses for social perception was also observed in the classification accuracies so that highest accuracy was observed in occipital and temporal areas, followed by parietal cortices and frontal and cingulate cortices. Lowest accuracies were found in the subcortical regions. (**Figure 5b**). We also validated that this gradient was not an artefact stemming from the sizes of the ROIs, as similar gradient was observed in the regional classification with ANOVA feature selection limited to 119 voxels (**Figure 5c**). **Figure SI-8** shows the statistical significance of the classification accuracies between all pairs of ROIs confirming the observed gradient in classification accuracies. Occipital and temporal areas (excluding Temporal pole) showed significantly higher classification accuracy than frontal and subcortical regions.

### Relationship between classification accuracy and ISC

Regional classification accuracy and ISC were positively correlated (Pearson r = 0.85, **Figure 6a**). Most occipital regions, STG and MTG showed high synchrony (ISC > 0.1) and high classification accuracy (acc > 0.25). Most parietal regions showed average ISC and average classification accuracy while frontal and subcortical regions showed low ISC and low classification accuracies. The most notable exception to this pattern was Heschl gyrus which had high ISC (0.28) yet average classification accuracy (acc = 0.22). **Figure 6b** summarizes the results from separate regression, ISC and classification analyses where the findings overlap most in temporal and occipital cortices.

**Figure 6.**
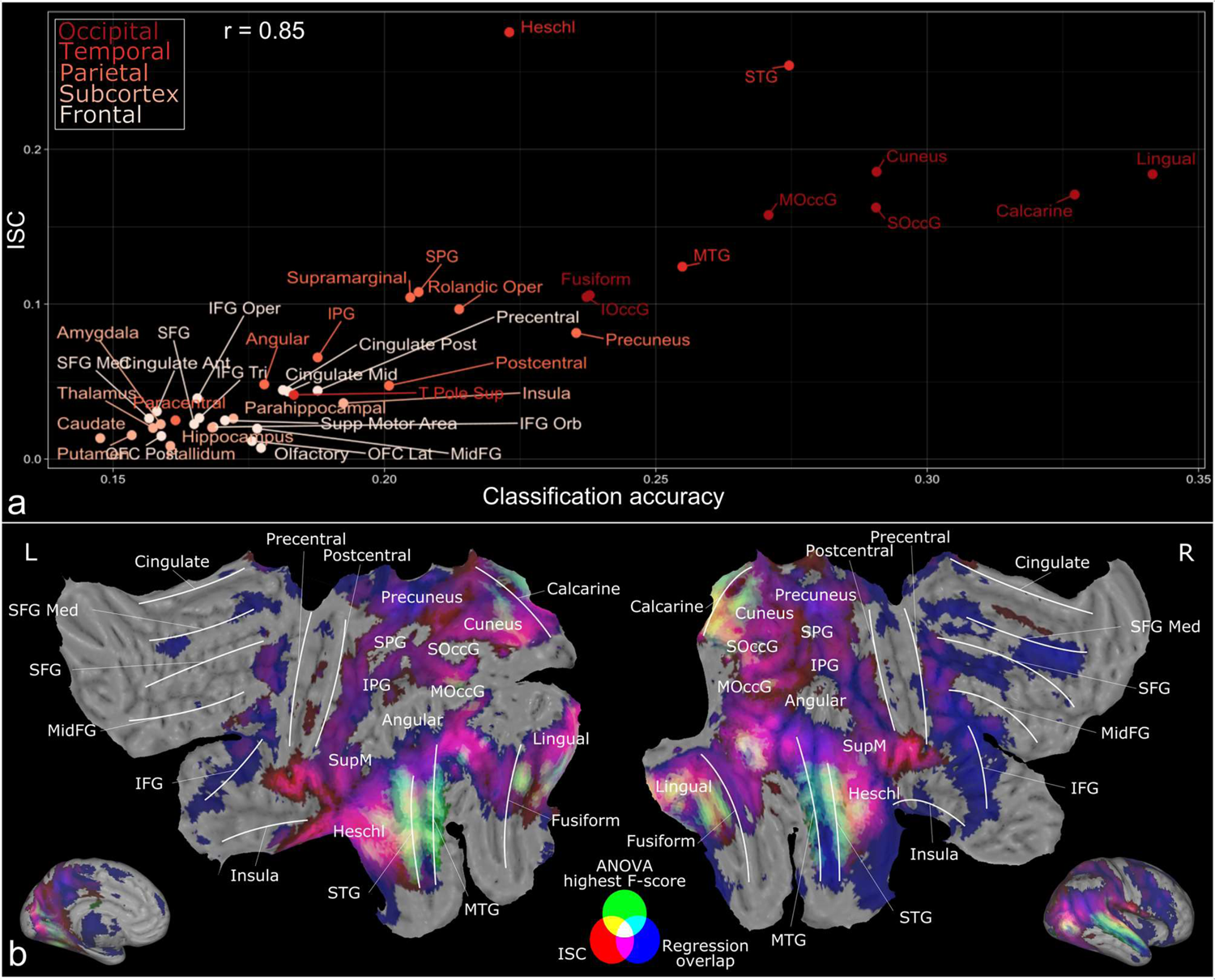
(a) Scatterplot showing the relationship between regional ISC and classification accuracy. (b) Additive RGB map summarizing the main findings. The overlap between activation patterns for perceptual dimensions in regression analysis is shown as blue (areas where at least 3 dimensions expressed FDR-corrected brain activation). Significant (FDR-corrected, q = 0.05) ISC across subjects is shown as red and the ANOVA selected voxels for the whole brain classification are shown as green.

## Discussion

Our findings provide the currently most detailed map of the social perceptual mechanisms in the human brain using naturalistic stimulus. The behavioral data established that 13 social dimensions reliably capture the social perceptual space contained in the video stimulus. The cerebral topography for social perception was organized along an axis, where posterior temporal and cortical regions served a central general-purpose role in social perception, while the regional selectivity for social dimensions increased towards frontal and subcortical regions. Multivariate pattern recognition established that particularly occipito-temporal and parietal regions carry detailed and spatially dimension-specific information regarding the social world, as evidenced by the highest classification accuracies in the multi-class classification approach. Both classification accuracy and consistency of the responses for specific social dimensions were the highest in the brain regions having most reliable (indicated by ISC) activation patterns throughout the experiment. These effects were observed although low-level sensory features were statistically controlled for. Altogether these results show that multiple brain regions are jointly involved in representing the social world and that different brain regions have variable specificity in their spatial response profiles towards social dimensions.

### Dimensions of social perception

The behavioural experiment established that the observers used consistently a set of 45 descriptors when evaluating the social contents of the movies. Dimension reduction techniques further revealed that these 45 features could be adequately summarized in 13 social dimensions. The largest clusters were organized along the valence dimension of the social interaction containing prosocial (e.g., kissing, touching, sexuality) versus antisocial (hurting others, yelling) behaviors. Social communicative behaviors (e.g., eye contact, talking) and body movements (e.g., waving, moving a foot) also formed large clusters. Play-related behaviors (laughing, playfulness) as well as feeding-related actions (e.g., tasting, eating) were also represented into smaller clusters. Notably, some features such as presence of males versus females, walking, and using objects remained independent of any of the clusters. Average hierarchical clustering algorithm was used because it yields clearly interpretable clusters and because feature similarity could be measured with correlation instead of absolute distance. Further research could establish how behavioural clusters found with hierarchical clustering relate to, for example, principal components off the same data and how the clusters generalize to other naturalistic stimuli.

The stimulus film clips cannot portray all possible social scenarios and Hollywood films are only a proxy of real-life social interaction. Still, 99 of the predefined 112 social features had sufficient occurrence rate in the stimulus video clips (**Figure SI-1**) which indicate that the stimulus contains a broad range of social information. The average duration of film clips was ~10 seconds and we acknowledge that this timescale does not allow examination of social processes occurring at slower temporal frequencies such as pair bonding and long-term impression formation. However, social perception may be astonishingly fast. Semantic, social, and affective categorization may happen in few hundred milliseconds (Nummenmaa et al., 2010) and the judgements do not significantly change from the initial judgments after longer consideration (Willis & Todorov, 2006). Electroencephalography (EEG) has also confirmed reliable associations between social perceptual features and brain response already 400ms after the stimulus (Dima et al., 2022) concluding that short video clips can capture some temporal scales of social perception. Data-driven models for characterising social perception (Adolphs et al., 2016) constitute an important, complementary alternative for the theory-based models for separate taxonomies of person, situation, and action perception since i) the found clusters are based on the actual perception of the social context, ii) the data-driven model does not separate persons, situations, and actions but is based on the subjects’ percept of the stimulus and iii) only dimensions actually present in the stimulus are considered. Importantly, this data-driven model for social perception has many similarities with previously proposed taxonomies. The largest observed clusters prosocial and antisocial behaviour closely relate to the emotional valence which is at the core of emotion theory (Russell, 1980) and is also considered in taxonomies describing persons (Simms, 2007) and situations (Parrigon et al., 2017; Rauthmann et al., 2014). Clusters play and feeding closely relate to dimensions Humor from situation taxonomy (Parrigon et al., 2017) and Food from action domain (Thornton & Tamir, 2022), respectively. Mapping of the neural space for social perception requires the social features to be consistently rated among the independent set of annotators. 61 of the total 112 rated social features showed low between-rater agreement (ICC < 0.5, **Figure SI-1**) which is itself an important finding regarding the consistency of the perceptual taxonomy individuals use for describing social events. The exclusion of these features had the effect that more abstract, or idiosyncratically judged dimensions cannot be addressed in this experiment and pushed the studied perceptual processes towards action and situation domains. Further research should nevertheless investigate the shared versus idiosyncratic social evaluations across individuals, as this would be informative regarding what are the core building blocks of the social environment that are shared across most observers.

### Cerebral gradient in social perception

The univariate BOLD-fMRI analysis based on social dimensions revealed that a widely distributed cortical and subcortical networks encode the social contents of the video stimuli. Most dimensions activated LOTC, STS, TPJ, as well as other occipitotemporal and parietal regions. There was a gradual change from these unselective social responses in occipitotemporal and parietal regions towards more selective responses in frontal and subcortical regions, suggesting that social perception is mainly processed in lateral and caudal parts of the brain. This effect was also confirmed by the ROI analysis. Most consistent responses were observed in all occipital regions and in temporal regions STG and MTG (which outline STS) and Heschl gyrus. In parietal cortex, most consistent responses were observed in supramarginal gyrus (a part of TPJ), SPG and precuneus. Frontally the responses were less consistent although brain activity in IFG, precentral gyrus and frontal part of medial SFG associated with a limited number of dimensions including “Prosocial behaviour”, “Antisocial behaviour”, “Feeding” and “Using an object”. These data are consistent with previous univariate studies addressing social functions for LOTC (Downing et al., 2001; Lingnau & Downing, 2015; Wurm & Caramazza, 2019; Wurm et al., 2017), STS (Deen et al., 2015; Isik et al., 2017; Lahnakoski et al., 2012; Walbrin et al., 2018), TPJ (Carter & Huettel, 2013; Saxe & Kanwisher, 2003), and MFC (de la Vega et al., 2016). The results were controlled with an extensive set of PCA rotated audiovisual features. A non-social regressor was also built from the stimulus time points where no social interaction was present, and this feature was added to the low-level model. The fMRI data were collected in one scan, hence ruling out the possibility to control for low-level features by cross-validation. Therefor the separation of social perceptual features from all possible low and mid-level features is not possible. However, we did not find extensive associations between social dimensions and BOLD responses in V1 and higher-level information such as body parts and actions have already been shown to associate with BOLD response better than low-level visual features in occipital cortex outside V1 (Tarhan & Konkle, 2020). Additionally, it has been shown that social features of actions explain more variance of EEG responses to videos than low-level visual features (Dima et al., 2022) further supporting the conclusion that the results reflect social information processing rather than low-level audiovisual perception.

### Decoding of perceptual social dimensions from brain activation patterns

The univariate analysis revealed the overall topography and regional brevity of the tuning for different social signals. However, this analysis cannot determine whether a single anatomical region activated by multiple social dimensions reflects responses to shared features across all the dimensions (such as biological motion perception or intentionality detection; Allison et al., 2000; Nummenmaa & Calder, 2009), or spatially overlapping yet dimension-specific processing. Multivariate classification analysis revealed that the answer to this question depends on the region. The ANOVA feature selection for the whole-brain classification retrieved voxels from STS, LOTC, TPJ and FG (**Figure 5** and **Figure 6**) yielding classification accuracy exceeding 50% for the multi-class classification. Regional classification confirmed that occipital, temporal and parietal regions showed average to high classification accuracies, whereas the classification accuracies diminished towards chance level in frontal and subcortical regions. These results show that even if the regional univariate responses for social dimensions were overlapping the specificity of the spatial activation patterns was different between regions. Interestingly, the ANOVA selected voxels found to best discriminate social features closely resemble the network proposed for social aspects of human actions in a recent study (Tarhan & Konkle, 2020) with the exception that our results are more bilateral. Previous multivariate studies have shown how individual social features are represented in these regions. For example, specific response patterns to pictures of faces versus animals, houses or man-made objects can be found in FG and LOTC (Haxby et al., 2001) and semantic information from different human actions judged from static images are represented in LOTC (Tucciarelli et al., 2019). Subsequent classification studies have shown that, for example, different facial expressions can be classified from activation patterns in FG, and STS (Said et al., 2010; Wegrzyn et al., 2015) and goal-oriented actions in LOTC and interior parietal lobe (Smirnov et al., 2017; Wurm & Lingnau, 2015). Importantly, our results show that BOLD-fMRI can be used for classification of multiple overlapping event categories from continuous naturalistic stimulation. Previous multivariate pattern analyses of social categories have used block designs and categorical stimuli matching the *a priori* category labels. In addition, these studies have only focused on a certain detailed aspect of socioemotional processing. The present results thus underline that even with high-dimensional naturalistic stimulus, the response properties of certain brain areas show high degree of category specificity.

The results from the classification analysis complement the results from the regression analysis with some limitations. The video clips were shown in the same order for all subjects, which may artificially boost classification accuracy, although the model should not learn the actual order of the events since the data were shuffled in each learning iteration. Regardless, the observed differences in classification accuracies in different brain regions should not be due to the order of the stimulus which is more interesting than the actual classification accuracies. It is likely that people focus attention in the most salient social details in the stimulus films instead of continuously monitoring for multiple sources of information with possibly low importance. Hence, we chose a classification approach where each time point was labelled with the social dimension of the highest relative intensity instead of trying to predict the values of all social features simultaneously. Future studies could try to predict multiple intensities for multiple categories in the stimulus set. Due to naturalistic and uncontrolled stimuli the classification dataset was unbalanced. Even in regions with near chance level total accuracy, some classes with large number of events were classified with relatively high accuracy (**Figure SI-9**) which may reflect the differences in the number of events in these classes and might not reflect the actual social information processing in the brain. Consequently, regional differences in the prediction accuracies to individual classes cannot be addressed.

### Reliability versus specificity of responses to social perceptual dimensions

We observed robust inter-subject correlation of brain activity in temporal and occipital regions while subjects viewed the video clips. Previous studies have found that the ISC is in general the strongest in sensory regions, and it progressively becomes weaker toward the polysensory and associative cortices (Hasson et al., 2010). Our data revealed that the strength of the ISC was contingent on the number of social features each region responded to in the univariate analysis (r = 0.86, **Figure 4c**). Additionally, regional ISC was also associated with the corresponding regional classification accuracy (r = 0.85, **Figure 6a**). These data highlight the relevance of the social domain to the cortical information processing, as the consistency of the regional neural responses was associated with the brevity of the tuning for social signals in each region. In other words, regions responding to multiple social signals also do so in a time-locked fashion across subjects, whereas the responses become more idiosyncratic as they become more selective. Importantly, this effect was not just an artefact of the consistency of sensory cortical responses to social signals but was also observed in higher-order associative areas including LOTC and STS.

### Functional organization of social perception in the human brain

The regional response profiles towards social signals can be summarized based on the combination of the regional response consistency (univariate regression analysis), the spatial response pattern specificity (MVPA) and the reliability of the BOLD signal across subjects (ISC). First, posterior temporal and occipital regions responded consistently to most social dimensions, while the presence of specific social dimensions could also be classified accurately from these regions. High classification accuracy suggests that these regions already hold dimension-specific and integrated information regarding the social world. Additionally, these regions responded consistently to the social stimuli (as indicated by high ISC) across subjects. LOTC, STS, TPJ, FG and occipital regions thus constitute the most fundamental hubs for social perception in the human brain and are likely involved in integration of the multisensory information and semantic representations regarding the social events (Allison et al., 2000; Lahnakoski et al., 2012).

Second, Heschl gyrus, the site of the auditory cortex (Da Costa et al., 2011) responded consistently to social dimensions but the classification accuracy was only moderate in that region while the ISC of the response was the highest of all regions. This suggests that Heschl gyrus processes domain-general social (most likely auditory) information but does not carry detailed information about the distinct social dimensions, as evidenced by the weak ability to classify specific dimensions from this region. Third, parietal regions especially precuneus, supramarginal gyrus and SPG showed consistent responses with numerous social dimensions and yet their ISC and classification accuracies were only moderate. Previously precuneus have been linked with attention and memory retrieval (Cavanna & Trimble, 2006), supramarginal gyrus with phonological (Hartwigsen et al., 2010) and visual (Stoeckel et al., 2009) processing of words and SPG in visuospatial processing and working memory (Koenigs et al., 2009). These parietal regions thus likely respond to some general features of the social signals or idiosyncratic brain states associated with social dimensions.

Frontal and subcortical regions responded only to a limited number of social dimensions, and classification accuracy and ISC were remained low. The regression analysis showed some consistency in IFG, precentral gyrus, the frontal part of the medial SFG, amygdala and thalamus, yet the classification accuracies remained low. MFC have previously been associated with higher-level social and affective inference such as linking social processing with decision making, affective processing and theory of mind (Amodio & Frith, 2006; de la Vega et al., 2016). But previous classification studies have not found specificity for responses to social perceptual dimensions in frontal cortex (Haxby et al., 2001; Oosterhof et al., 2012; Wegrzyn et al., 2015; Wurm & Lingnau, 2015). Thus, frontal areas may subserve higher-order social process by linking low-level social perception into more complex and abstract cognitive processes such as making predictions of the next actions or linking perception with the brains affective system. Indeed, there is evidence that MFC could be responsible in giving a affective meaning for the ongoing experiences and that MFC processing is highly idiosyncratic (Chang et al., 2021). Finally, limbic regions such as amygdala and thalamus in turn have been linked with processing of (negative) emotions (Karjalainen et al., 2019) and accordingly they showed reliable responses primarily to perception of antisocial behaviours.

## Conclusion

Using a combination of data-driven approaches and multivariate pattern recognition we established the perceptual space for social features and mapped the cerebral topography of social perception that can be adequately described with 13 perceptual dimensions. Social perceptual space included clusters of social features describing prosocial and antisocial behaviour, feeding, body movement, communication and playfulness, as well as individual dimensions male, female, running, walking, searching, crying and using an object. Clear gradient in response selectivity was observed from broad response profiles in temporal, occipital and parietal regions towards narrow and selective responses in frontal and subcortical regions. Perceptual social dimensions could be reliably decoded from regional activation patterns using multivariate pattern analysis. Both regression analysis and multivariate pattern analysis highlighted the importance of LOTC, STS, TPJ and FG and other occipitotemporal regions as dimension-specific social information processors, while parietal areas and Heschl’s gyrus process domain-general information from the social scenes. Additionally, regional response profiles for social perception closely related to the overall reliability of the BOLD responses. Altogether these results highlight the distributed nature of social processing in the brain as well as the spatial specificity of brain regions to social dimensions.

## Acknowledgements

The study was supported by grants to LN from European Research Council (#313000) and the Academy of Finland (#332225, #294897). The authors declare no competing financial or non-financial interests. We thank Tuulia Malen for the fruitful conversations around the analysis methods used in this article and Juha Lahnakoski for the help with low-level feature extraction.

## Author contributions

SS developed the analysis methods, analysed the data, and wrote the manuscript.

TK preprocessed the fMRI data, developed the regression analysis methods, and wrote and reviewed the manuscript.

SNF developed the MVPA analysis methods and reviewed the manuscript.

MH conceptualized the study design, collected the data, and reviewed the manuscript.

VP, KS, and LS collected the data and reviewed the manuscript.

EG developed the analysis and preprocessing methods and reviewed the manuscript.

JH reviewed the MR images and reviewed the manuscript and HK supervised the security of MRI data collection and reviewed the manuscript.

LN conceptualized the study design, acquired funding, supervised the project, developed analysis methods, and wrote and reviewed the manuscript.

## Data availability

The stimulus movie clips can be made available for review although copyrights preclude public redistribution of the stimulus set. Short descriptions of each film clip can be found from a supplementary excel sheet. According to Finnish legislation, the original (even anonymized) neuroimaging data used in the experiment cannot be released for public use. All necessary files can however be made available for reviewers. The brain activation patterns (unthresholded T-maps) for each GLM predictor are available in NeuroVault (https://neurovault.org/collections/IZWVFEYI/).

## Code availability

We developed in-house scripts for low-level audiovisual feature extraction and ridge regression optimization, and the scripts are freely available in GitHub (https://github.com/santavis/functional-organization-of-social-perception). Other analyses involved using available R and Python packages and these scripts can be made available for review.

## Footnotes

1 Dimensions “Male” and “Female” were excluded from classification, because unlike the rest of the dimensions, they are genuinely binary features and thus not comparable with the other dimensions in the implemented classification framework (see **Figure SI-4**).

