## Supplementary Materials for "Functional organization of social perception in the human brain"

#### Supplementary methods

**Hierarchical clustering of social features**

Unweighted pair group method with arithmetic mean (UPGMA), as implemented in R, was used as the clustering algorithm (https://www.rdocumentation.org/packages/stats/versions/3.6.2/topics/hclust). Other average linkage clustering methods implemented in the R package (WPGMA, WPGMC and UPGMC) yielded highly similar clustering hierarchy. Hierarchical clustering requires a desired number of resulting clusters as an input for automatic definition of cluster boundaries from hierarchical tree (**Figure SI-2**). To estimate the optimal number of clusters we chose three criteria that the clustering result should satisfy. These were cluster stability, theoretically meaningful clustering, and sufficient reduction in collinearity between the clusters. To assess the stability of clusters with different number of clusters we conducted a consensus clustering analysis with ConsensusClusterPlus R package (Wilkerson & Hayes, 2010). In consensus clustering analysis 80% of stimulus time points and 80% of social features were randomly sampled and then clustered using UPGMA. Data were resampled 5000 times and then the stability of clusters with different number of clusters (*k*) was assessed based on the consensus of the resampling (**Figure SI-3**). Analysis showed that clustering results stabilize when $k > 10$ (Relative change in area under curve between CDFs of the consensus matrix of *k* clusters and $k$ - 1 clusters is minimal, **Figure S1-3 a & b**). In addition, $k = 13$ was the lowest number of clusters yielding in theoretically meaningful cluster labels. With 13 clusters, maximum pairwise correlation between any two resulting regressors of the model was below 0.4 and maximum VIF value was 3.3 (male regressor) which was considered as sufficient reduction in collinearity.

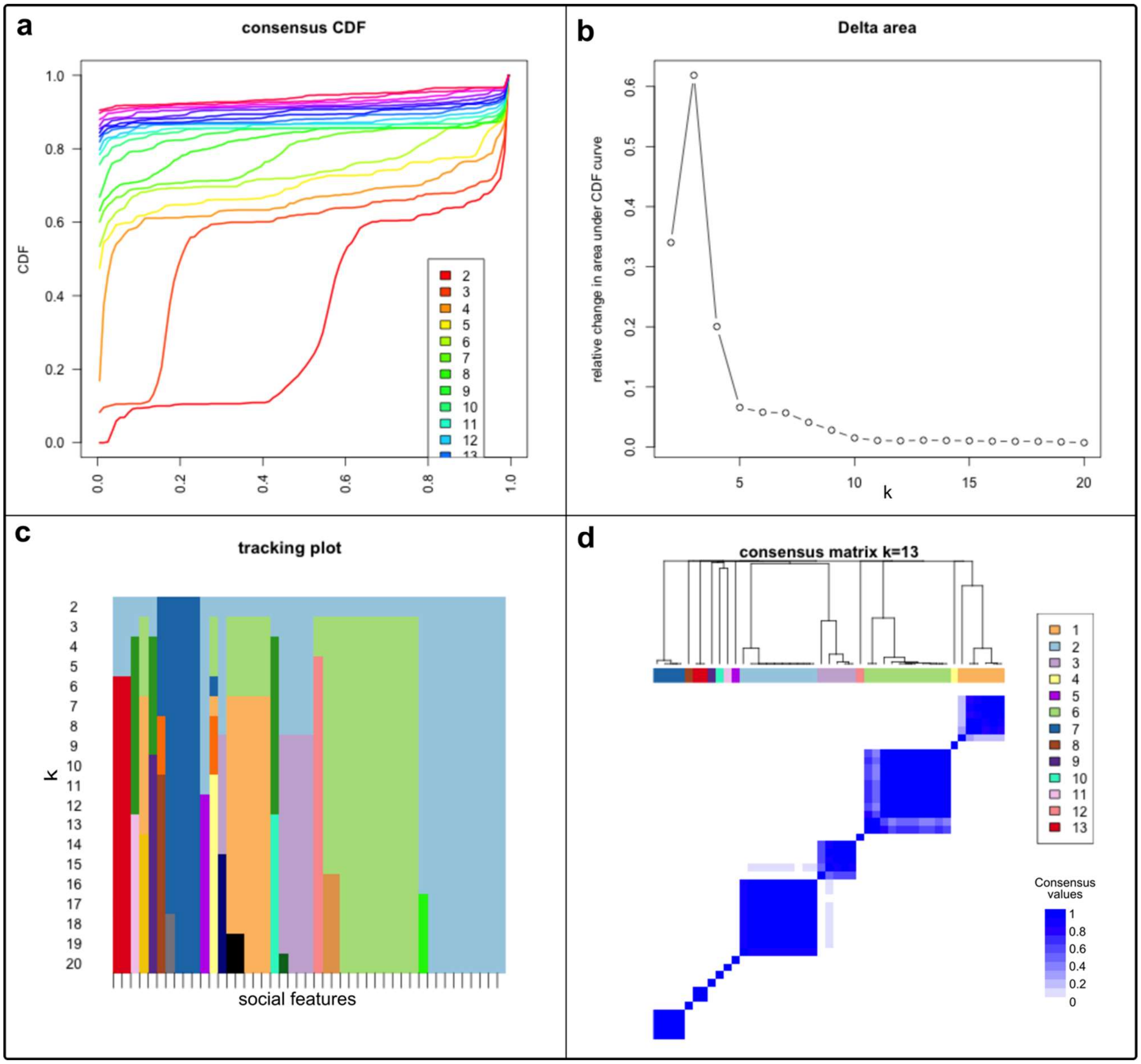


***Supplementary Figure 3.*** *Results of the consensus clustering analysis with UPGMA as the clustering algorithm. a) Cumulative distribution function (CDF) of the consensus matrix for different number (k) of clusters. b) Relative change in area under curve between CDF of k clusters and k-1 clusters. c) Tracking plot showing changes in cluster structures with different number of clusters. d) Heatmap of the consensus values (1=always clustered together, 0=never clustered together) between social features (k=13).*

### Ridge regression

Ridge regression uses L2-regularization where OLS β-coefficient estimators are penalized in following formula:

$$\hat{\beta} = (X^T X + \lambda I)^{-1} X^T y ,$$

where λ is the ridge parameter which controls the amount of bias induced into the model. The ridge parameter was optimised using leave-one-subject-out cross-validation. β-coefficients were estimated over N-1 subjects and then the BOLD signal of remaining subject was predicted with estimated β-coefficients. Prediction error (PE) for the remaining subject was calculated as:

$$PE = \sqrt{\sum_{1}^{K} (y - \hat{\beta}\, X)^2},$$

where K is the number of voxels and y is the BOLD signal in the i:th voxel. Cross-validation was repeated for every subject and average prediction error over all subjects was used as a measure of the model's fit to the data. Automatic optimiser function (https://www.mathworks.com/help/matlab/ref/fminbnd.html) was used to optimise ridge parameter for minimum prediction error and then β-coefficients for all subjects were estimated with optimised ridge parameter (λ = 83 for initial low-level regression and λ = 28 for the following social + low-level model). Prior to statistical modelling the BOLD signals were divided by their mean to make the regression coefficients more comparable between different individuals (Chen et al., 2017). A flowchart of ridge regression optimisation process is shown in **SI-Figure 6**.

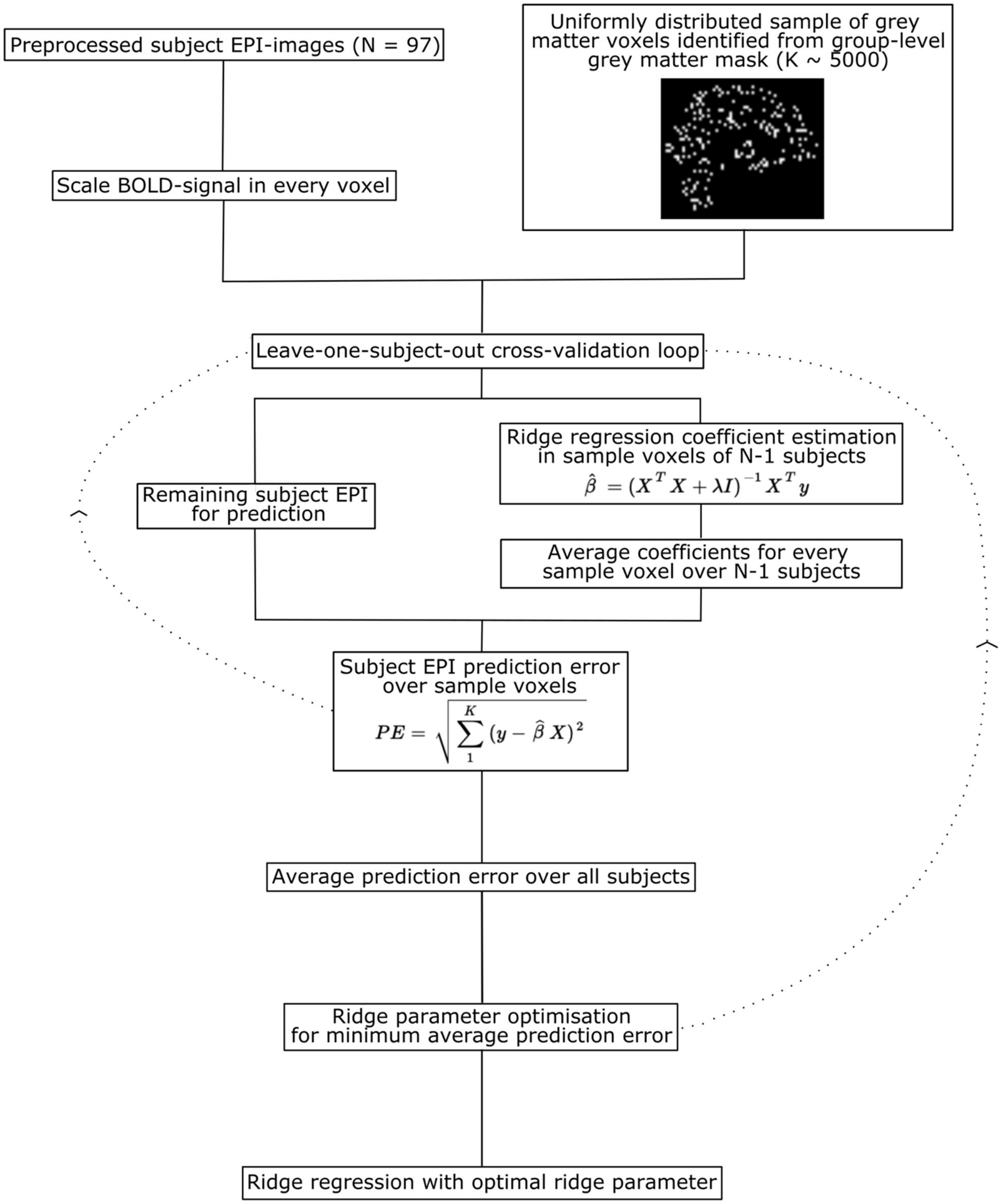


***Supplementary Figure 6***. *Ridge regression optimisation process. Ridge regression parameter (λ) was optimised using leave-one-subject-out cross-validation.*

#### Comparison of low-level and social models

In addition of controlling low-level features in the analyses for social dimensions we also compared the predictions of separate low-level and social models as supplementary analysis. First, we run separate Ridge regression analyses with the low-level model (11 predictors) and with the social model (13 predictors), where no low-level features were controlled. Then, we compared the predictions of the models by calculating predictive $R^2$-values for each subject in the leave-one-subject-out cross validation process and then averaging the voxelwise predictive $R^2$-values over all subjects. Because the models had unequal number of predictors, we used adjusted $R^2$ for the model comparison. In the results we report where in the brain the social model gave more accurate predictions than the low-level model in terms of predictive adjusted $R^2$ (FDR-corrected, q=0.05). We also run the classification analysis only in these voxels to assess the classification accuracy of these regions compared to the whole brain or anatomical ROI classification accuracies.

### Supplementary Results

**All annotated socioemotional features, N = 112**

| Sensory input | Basic body functions | Person characteristics | Inner states | Social interaction signals | Social interaction characteristics |
|---|---|---|---|---|---|
| Smelling | Hand-movement | Age | Thinking | Communicating with gestures | Friendship |
| Pain | Turning head | Distance form other people | Dreaming | Singing | Equality |
| Looking | Turning body posture | Pleasantness | Feeling uncertain | Dancing | Inequality |
| Tasting | Jumping | Unpleasantness | Confidence | Talking | Attachment |
| Sensing touch | Falling | Attractiveness | Empathy | Listening | Intimacy |
| | Lifting something | Kindness | Self-control | Laughing | Romanticism |
| | Facial expressions | Trustworthiness | Emotional warmth | Crying | Formality |
| | Other facial movement | Competence | Remembering | Yelling | Informality |
| | Eye-blinking | Dominance | Feeling of success | Moaning | Personal |
| | Gaze orientation | Nakedness | Feeling secure | Kissing and hugging | Superficiality |
| | Coughing and snotting | Male | Thirst | Eye-contact | Compliance |
| | Touching an object | Female | Being tired | Touching a human | Using authority |
| | Sleeping | | Disappointment | Hurting others | Voluntarity |
| | Drinking | | Imagining | | Cooperation |
| | Counting | | Wanting | | Reluctance |
| | Reading | | Being drunk | | Violence |
| | Writing | | Calmness | | Morality |
| | Body-movement | | Making a desicion | | Conflict |
| | Foot-movement | | Observing | | Number of people |
| | Running | | Satisfaction | | Human voice |
| | Walking | | Social cohesion | | Sexuality |
| | Lying (lay) | | Loneliness | | Bodiliness |
| | Standing | | Goal orientation | | Playfulness |
| | Waving | | Pleasant feelings | | Hostility |
| | Sitting | | Unpleasant feelings | | Aggression |
| | Using an object | | Hunger | | |
| | Searching | | Passion | | |
| | Eating | | Arousal | | |
| | | | Feeling of sickness | | |

**Data-driven clusters of selected socioemotional features, N = 13**

| Prosocial behaviour | Antisocial behaviour | Communication | Body-movement | Feeding | Play | Individual features |
|---|---|---|---|---|---|---|
| Kissing and Hugging | Pain | Talking | Body-movement | Hunger | Laughing | Male |
| Pleasant feelings | Aggression | Listening | Foot-movement | Eating | Playfulness | Female |
| Passion | Conflict | Human voice | Dancing | Tasting | | Walking |
| Sexuality | Hostility | Eye-contact | Waving | Sitting | | Running |
| Nakedness | Feeling of sickness | Looking | Number of people | | | Searching |
| Bodiliness | Morality | Standing | | | | Crying |
| Sensing touch | Hurting others | | | | | Using an object |
| Touching a human | Violence | | | | | |
| Moaning | Unpleasant feelings | | | | | |
| Lying (lay) | Yelling | | | | | |
| | Arousal | | | | | |

***Supplementary Table 1****. Complete listing of annotated socioemotional features (N=112). Features with insufficient inter-rater reliability or occurrence rate are marked with red color. Data-driven clusters (perceptual dimensions) of selected features (N = 13) are shown in the bottom section.*

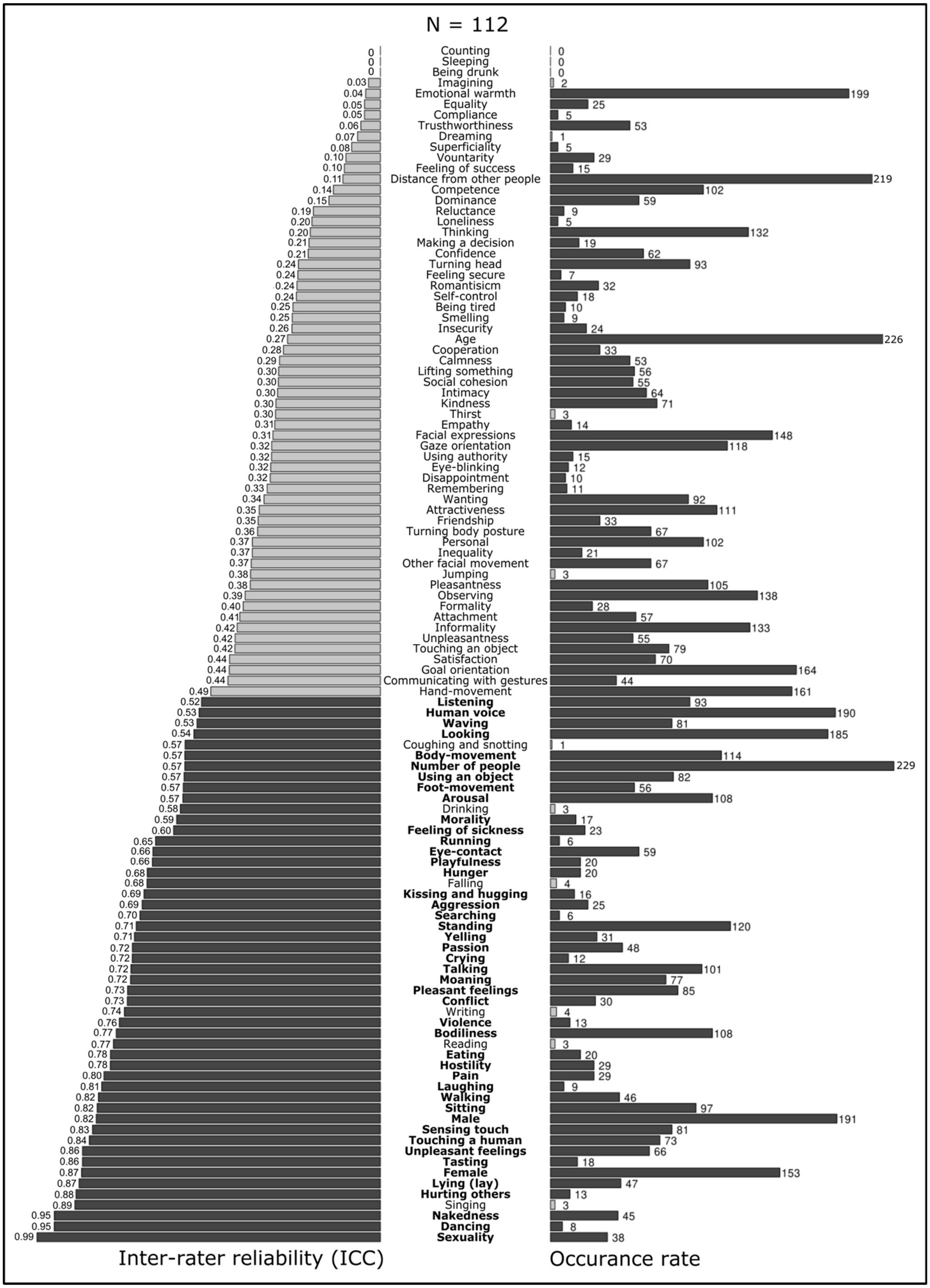

N = 112
Inter-rater reliability (ICC)
Occurance rate

***Supplementary Figure 1.*** *Inter-rater reliability and occurrence rate for all 112 features. Selected features (N=45) are bolded.*

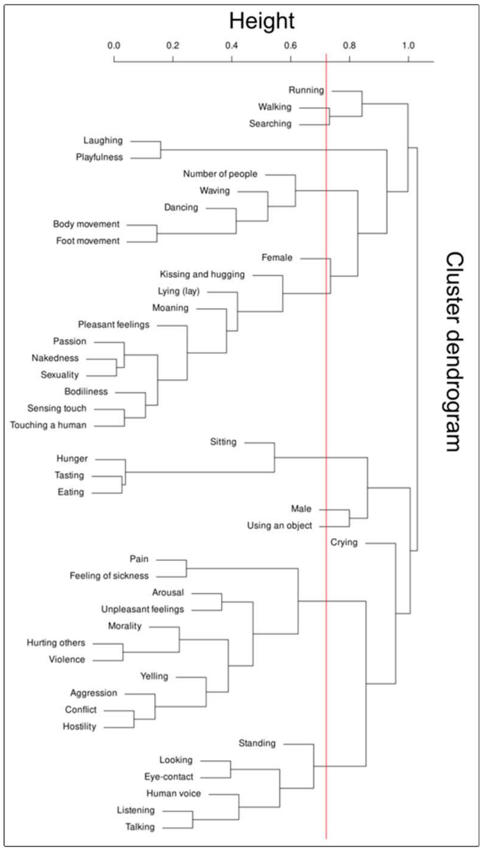


***Supplementary Figure 2.*** *Hierarchical dendrogram of selected features and a vertical line indicating height of cluster boundaries.*

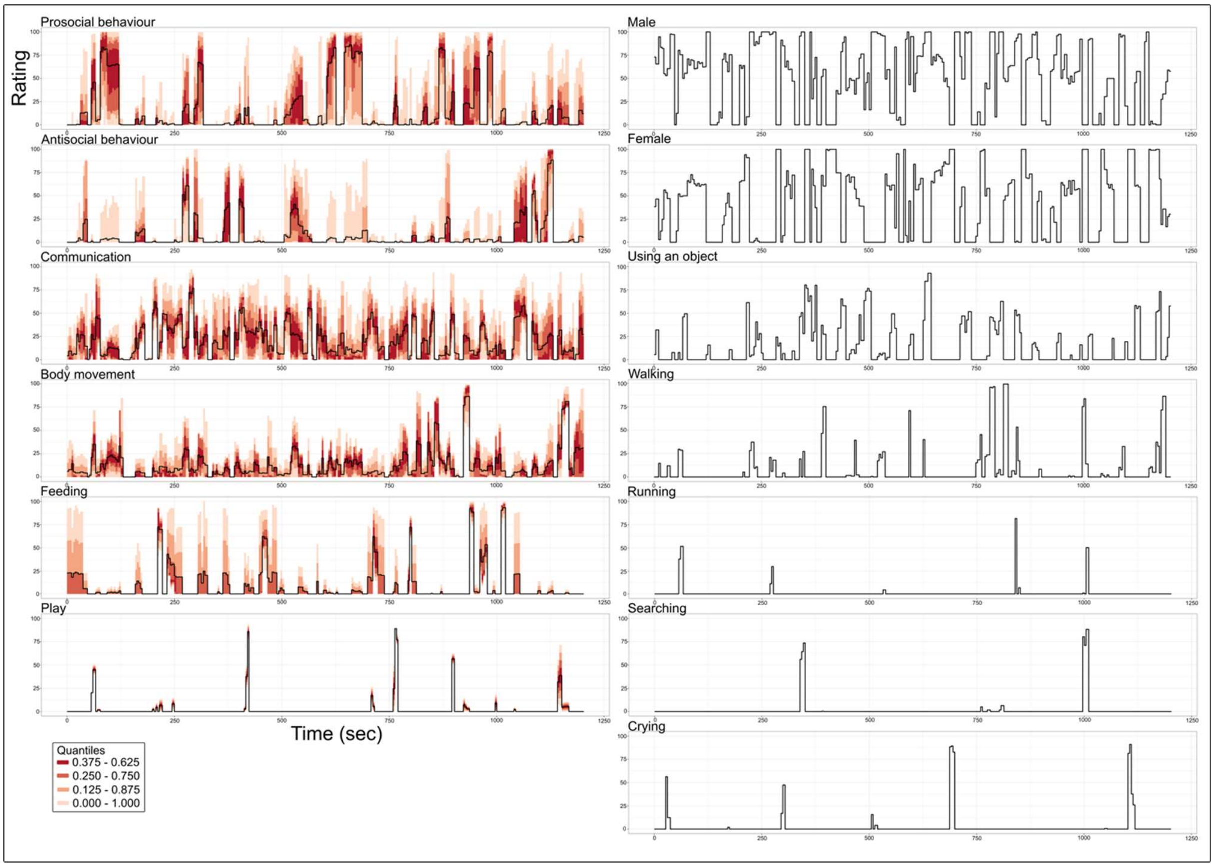


***Supplementary Figure 4.*** *Feature rating time series for each social dimension. The time series for cluster dimensions show the mean rating of all social features within the cluster. Quantiles of individual feature ratings are visualized around the average time series of each cluster.*

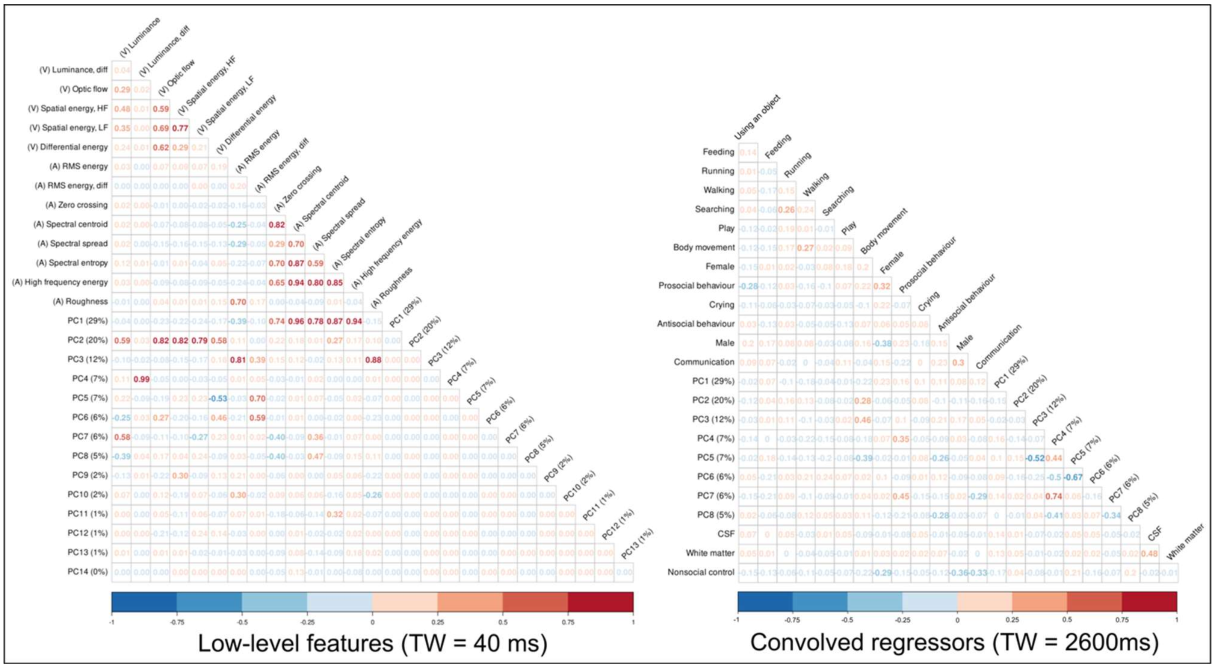


***Supplementary Figure 5.*** *Correlations between low-level audiovisual features and main social dimensions. The left panel shows the extracted low-level audiovisual features and their correlations with corresponding principal components (PCs) in unconvolved form. The right panel shows the correlation between convolved regressors.*

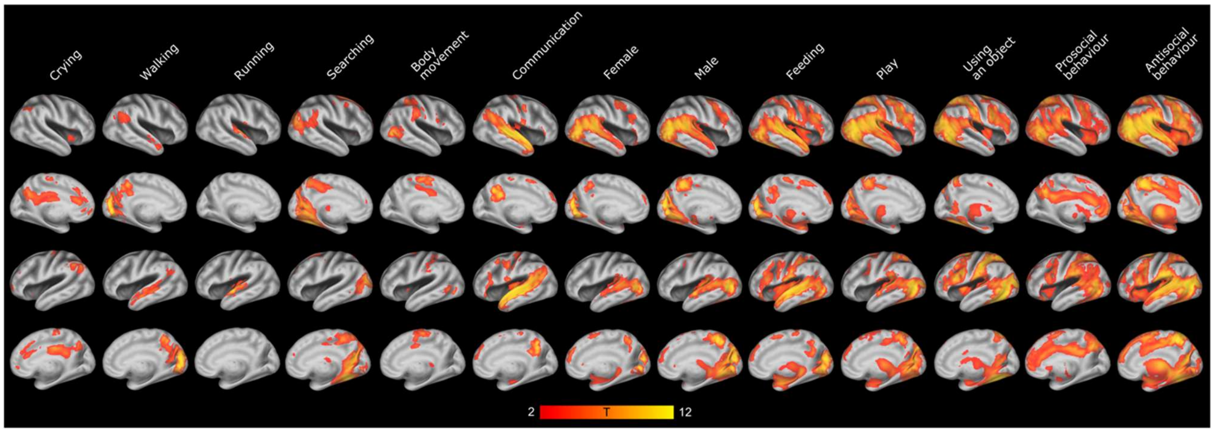


***Supplementary Figure 7.*** *Brain regions showing increased BOLD activity for the social dimensions. Results show the voxelwise T-values (FDR-corrected, q = 0.05) of increased BOLD activity for each social dimension from the multiple regression analysis. See also Figure 2.*

| Total accuracy | Minimum accuracy (class) | Minimum precision (class) | SEM of total prediction accuracy | Param: hidden_layers | Param: nodes | Param: alpha | Param: max_iter | Runtime (min) |
|---|---|---|---|---|---|---|---|---|
| 0,482 | 0,415 | 0,424 | 0,0087 | 1 | 100 | 0,0001 | 500 | 73 |
| 0,473 | 0,404 | 0,416 | 0,0090 | 1 | 100 | 0,0001 | 1000 | 61 |
| 0,475 | 0,377 | 0,435 | 0,0086 | 1 | 100 | 0,1000 | 500 | 57 |
| 0,471 | 0,388 | 0,430 | 0,0086 | 1 | 100 | 0,1000 | 1000 | 58 |
| 0,486 | 0,409 | 0,442 | 0,0089 | 1 | 100 | 1,0000 | 500 | 57 |
| 0,483 | 0,410 | 0,437 | 0,0089 | 1 | 100 | 1,0000 | 1000 | 57 |
| 0,483 | 0,399 | 0,429 | 0,0083 | 1 | 200 | 0,0001 | 500 | 77 |
| 0,492 | 0,412 | 0,436 | 0,0087 | 1 | 200 | 0,0001 | 1000 | 78 |
| 0,486 | 0,399 | 0,423 | 0,0090 | 1 | 200 | 0,1000 | 500 | 75 |
| 0,488 | 0,410 | 0,414 | 0,0084 | 1 | 200 | 0,1000 | 1000 | 75 |
| 0,485 | 0,404 | 0,432 | 0,0088 | 1 | 200 | 1,0000 | 500 | 79 |
| 0,492 | 0,404 | 0,430 | 0,0085 | 1 | 200 | 1,0000 | 1000 | 78 |
| 0,501 | 0,436 | 0,450 | 0,0102 | 2 | 100 | 0,0001 | 500 | 58 |
| 0,502 | 0,407 | 0,453 | 0,0095 | 2 | 100 | 0,0001 | 1000 | 57 |
| 0,503 | 0,423 | 0,457 | 0,0089 | 2 | 100 | 0,1000 | 500 | 57 |
| 0,507 | 0,436 | 0,446 | 0,0095 | 2 | 100 | 0,1000 | 1000 | 58 |
| **0,519** | **0,451** | **0,476** | **0,0098** | **2** | **100** | **1,0000** | **500** | **59** |
| 0,519 | 0,454 | 0,460 | 0,0104 | 2 | 100 | 1,0000 | 1000 | 59 |
| 0,522 | 0,437 | 0,456 | 0,0094 | 2 | 200 | 0,0001 | 500 | 81 |
| 0,528 | 0,474 | 0,483 | 0,0095 | 2 | 200 | 0,0001 | 1000 | 81 |
| 0,528 | 0,445 | 0,469 | 0,0091 | 2 | 200 | 0,1000 | 500 | 76 |
| 0,524 | 0,432 | 0,473 | 0,0097 | 2 | 200 | 0,1000 | 1000 | 77 |
| 0,537 | 0,445 | 0,503 | 0,0099 | 2 | 200 | 1,0000 | 500 | 91 |
| 0,534 | 0,472 | 0,484 | 0,0101 | 2 | 200 | 1,0000 | 1000 | 91 |

***Supplementary Table 2.*** *Results of neural network classifier parameter tuning. The whole brain classification was performed with a set of different parameter values for the classifier algorithm. Optimal parameters (hidden layers = 2, nodes = 100, alpha = 1.0, maximum iteration = 500) for the final analysis was chosen based on the algorithm's prediction accuracies, precisions, and algorithm runtime.*

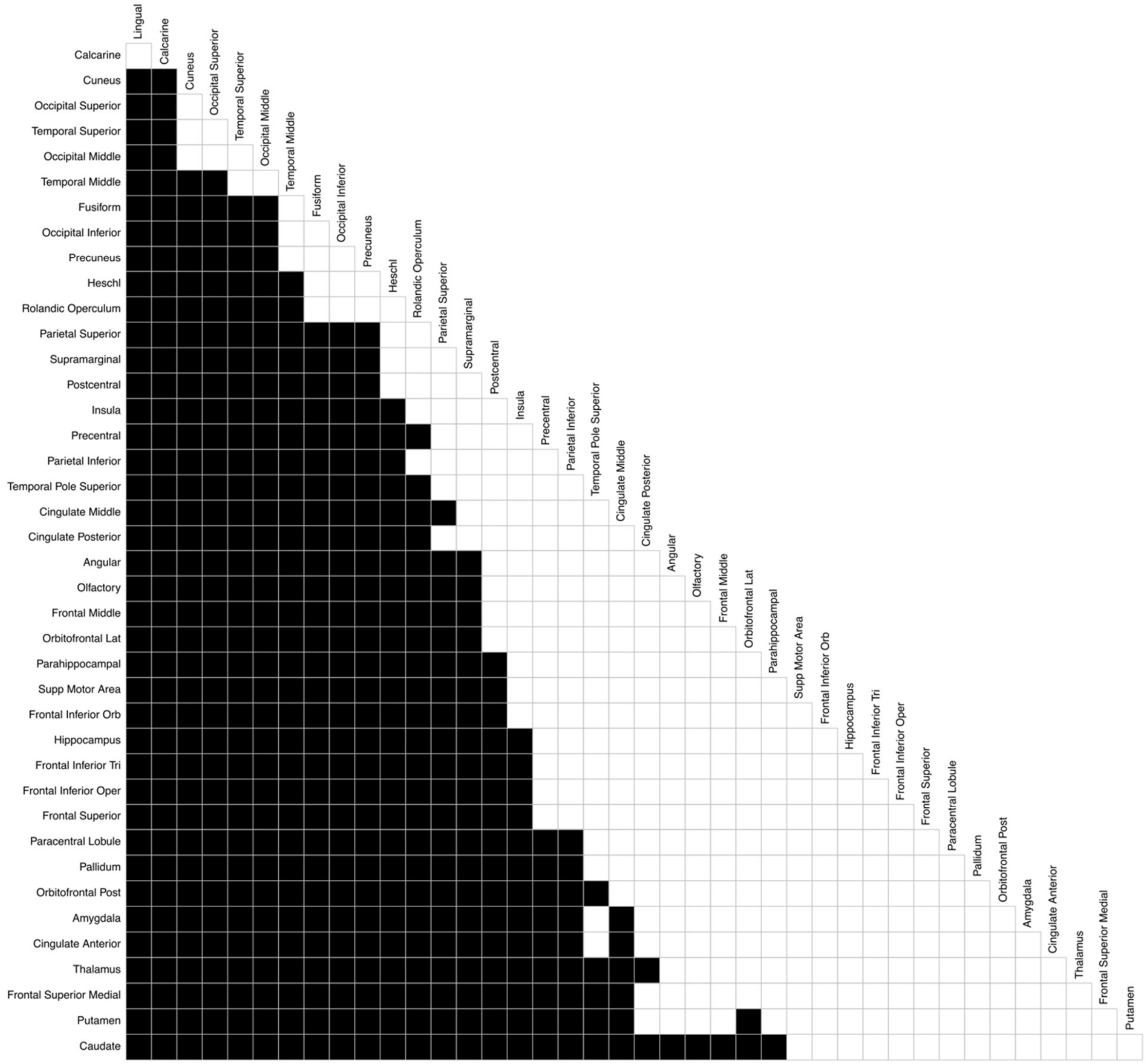


***Supplementary Figure 8.*** *Comparison of ROIs in classification accuracies for social dimensions. Significant (p<0.05, Bonferroni corrected, paired t-test) differences between ROIs are marked black and ROIs are ordered from the highest classification accuracy (Lingual) to the lowest (Caudate) similarly as in figure 5. Black rectangle states that the ROI in the column has significantly higher classification accuracy than the ROI in the corresponding row. Figure 5 visualizes the gradient in classification accuracies of social dimensions and this figure confirms the observed gradient statistically.*

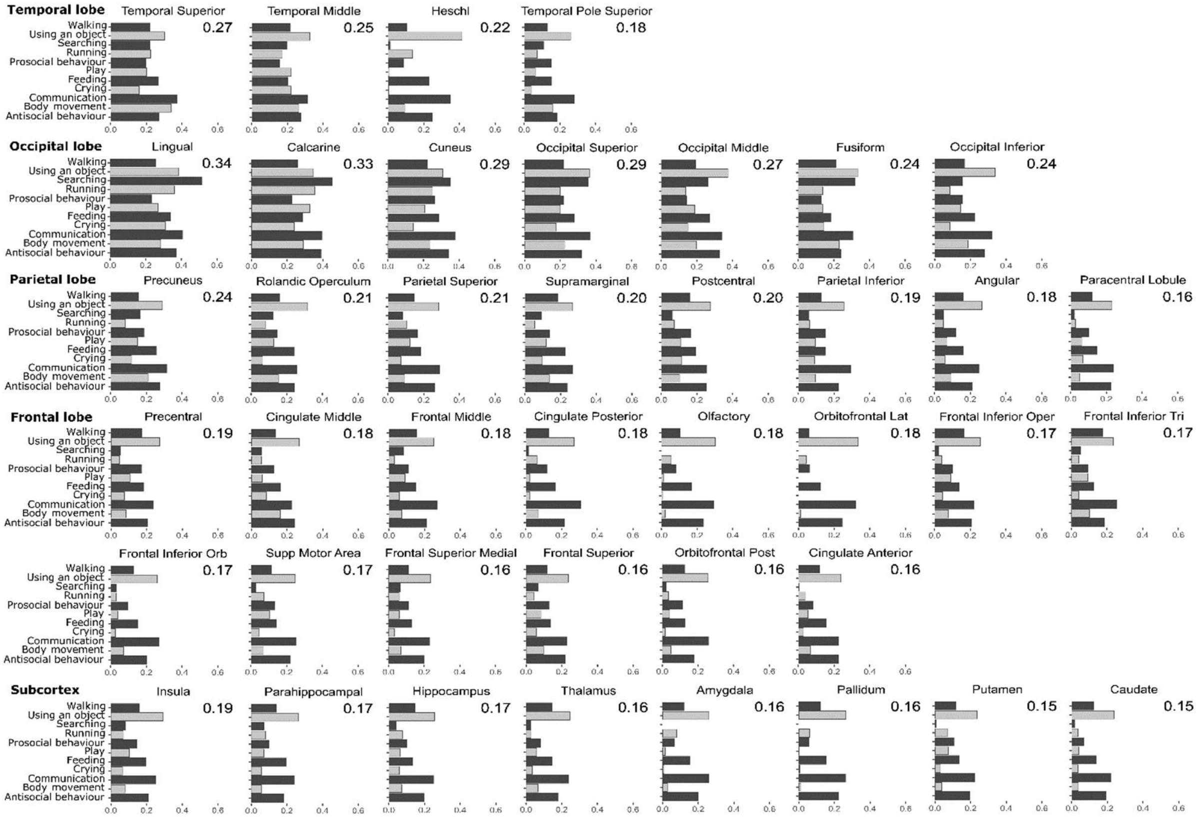


***Supplementary Figure 9.*** *Region-of-interest classification results for perceptual social dimensions. Regional differences in the prediction accuracies to individual classes cannot be addressed.*

### Comparison of low-level and social models

Separate Ridge regression analyses for BOLD signal were conducted for both low-level and social models. In terms of predictive adjusted $R^2$, the social model predicted the BOLD signal significantly better in many cortical areas including voxels from functional areas STS, LOTC, TPJ and anatomical areas FG, SPG, IFG and precentral gyrus (**Figure SI-9**). Low-level model including audiovisual properties and the mean signals from CSF and WM predicted the BOLD response significantly better in most brain regions including primary visual and auditory areas. Classification analysis using only voxels with significantly higher predictive adjusted $R^2$ compared to low-level model yielded in 35% ($p<0.01$) total classification accuracy which is comparable with the highest observed classification accuracy in anatomical ROIs (34% in lingual gyrus).

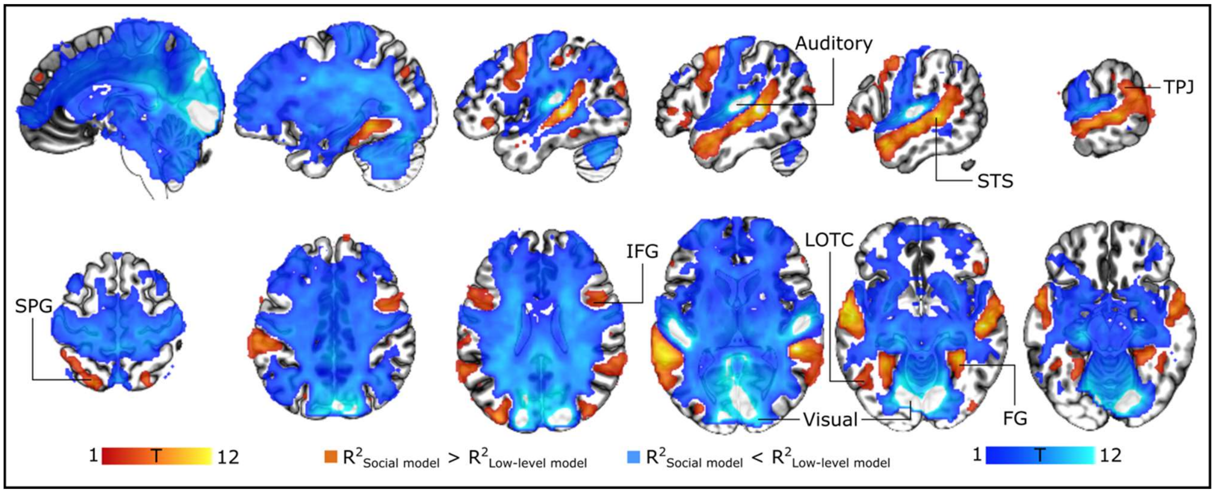


***Supplementary Figure 10.*** *Comparison of social and low-level models. Statistically significant (FDR corrected, q = 0.05) differences in predictive adjusted* $R^2$ *values between social and low-level models from multivariate regression analysis.*

| | |
|---|---|
| **Anatomical** | |
| Cingulate Ant | Anterior cingulate cortex |
| Cingulate Post | Posterior cingulate cortex |
| FG | Fusiform gyrus |
| IFG | Inferior frontal gyrus |
| IFG Oper | Opercular part of inferior frontal gyrus |
| IFG Orb | Orbital part of inferior frontal gyrus |
| IFG Tri | Triangular part of frontal inferior gyrus |
| IOccG | Inferior occipital gyrus |
| IPG | Inferior parietal gyrus |
| MidFG | Middle frontal gyrus |
| MOccG | Middle occipital gyrus |
| MTG | Middle temporal gyrus |
| OFC Lat | Lateral orbitofrontal cortex |
| OFC Post | Posterior orbitofrontal cortex |
| Olfactory | Olfactory bulb |
| Rolandic Oper | Rolandic operculum |
| SFG | Superior frontal gyrus |
| SFG Med | Medial superior frontal gyrus |
| SOccG | Superior occipital gyrus |
| SPG | Superior parietal gyrus |
| STG | Superior temporal gyrus |
| SupM | Supramarginal gyrus |
| Supp Motor Area | Supplementary motor area |
| T Pole Sup | Superior temporal pole |
| **Functional** | |
| aSTS | Anterior superior temporal sulcus |
| Auditory | Primary auditory cortex |
| LOTC | Lateral occipitotemporal cortex |
| MFC | Medial frontal cortex |
| pSTS | Posterior superior temporal sulcus |
| STS | Superior temporal sulcus |
| TPJ | Temporoparietal junction |
| Visual | Primary visual cortex |

***Supplementary Table 3.*** *Table of abbreviations used in this article. Anatomical abbreviations refer to AAL2 atlas-based regions-of-interest. Functional regions-of-interest are used when interpreting the results in comparison with previous literature.*
